# COMPARATIVE ANALYSIS OF THE RNA-CHROMATIN INTERACTOME DATA

**DOI:** 10.1101/2025.08.16.668492

**Authors:** G.K. Ryabykh, A.I. Nikolskaya, L D. Garkul, A.A. Mironov

## Abstract

Two types of experiments are used to study RNA-chromatin interactions: a search for the interactome of individual RNAs («one-to-all» or OTA) and a genome-wide search for contacts of all RNAs («all-to-all» or ATA). The article presents a comparative analysis of these data. The concept of «chromatin potential» is introduced, which shows the overrepresentation of contacts in ATA data compared to the expected value obtained from RNA sequencing data. It is shown that the use of this characteristic allows one to select RNAs that interact with chromatin more specifically. The consistency of the replicas is studied, and data from experiments of different types are compared. As a result of the analysis, estimates of the completeness and specificity of the contacts found are obtained.

## INTRODUCTION

Every year, more and more evidence appears that non-coding RNAs (ncRNAs) in animals and plants are involved in a wide range of biological processes, such as regulation of cell differentiation and gene expression, chromatin remodeling, maintenance of chromatin structure, splicing, RNA processing, formation of biomolecular condensates, and others. In addition, disruption of regulatory pathways mediated by ncRNAs can lead to various diseases [1].

RNA molecules interact with a variety of proteins, chromatin, and other RNAs. Experimental methods that allow identification of DNA loci that ncRNAs contact can be divided into two groups: «one-to-all» («OTA») and «all-to-all» («ATA»). The first group of methods (RAP [2], CHART-seq [3], ChIRP-seq [4], dChIRP-seq [5], ChOP-seq [6], CHIRT-seq [7]) determines contacts of previously known RNA with chromatin, and the second group of methods (MARGI [8], GRID-seq [9], ChAR-seq [10-11], iMARGI [12], RADICL-seq [13], Red-C [14]) is aimed at determining all possible RNA-DNA contacts in the cell [15].

The data from ATA experiments have a number of biases. First, there is an increased density of RNA contacts with chromatin near the gene encoding this RNA. This bias in the data will be called RD scaling. Second, the availability of chromatin, which we will call the background, has a significant impact. To estimate the background, either the «input» data in OTA experiments or mRNA contacts are used in ATA experiments [9]. In addition, by design, these experiments have limited accuracy of contact determination. In ATA experiments, cross-linking of RNA with chromatin can occur at some distance from the real contact (Figure 1a). At the same time, in OTA experiments, the accuracy of contact position determination depends only on the sizes of DNA fragments (Figure 1b). A special problem is the presence of non-specific interactions. A significant part of the observed contacts can be explained by electrostatic attraction between negatively charged RNA and positively charged histone tails, as well as by the preferential cross-linking of amino groups by formaldehyde [16] present on lysines and arginines of histones. Although the affinity of such non-specific interactions is relatively low, their combined contribution is significant due to the huge number of potential binding sites (Figure 1c). On the other hand, technical limitations of existing experimental methods lead to the loss of some of the true contacts. Therefore, the question arises about the specificity and completeness of the RNA-chromatin interactome data.

**Figure 1.**
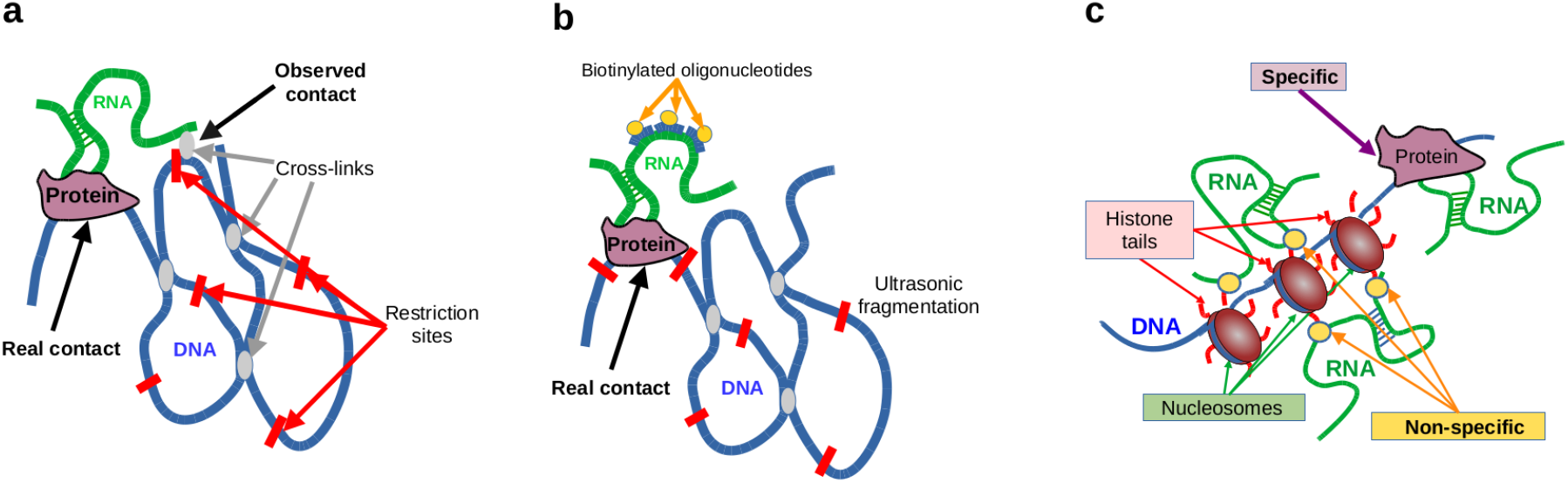
The accuracy of determining the position of the real contact differs in the ATA and OTA protocols. **a**: the source of the position shift in ATA data is the chromatin structure. **b**: the OTA shift of the observed contact position from the real one is determined only by the size of DNA fragments. **c**: a possible source of non-specific interactions.

The aim of this work is a comparative analysis of experimental data on the RNA-chromatin interactome to assess the accuracy, completeness and specificity of experimental data on RNA-chromatin contacts.

## MATERIALS AND METHODS

### Data

Human and mouse RNA-chromatin interactome data were retrieved from the RNA-Chrom database [17]. Of all ATA data, we retained only those for which RNA-seq data from the same cell line were found. If there were more than two replicates in the ATA data, we selected the two more complete replicates. RNA-seq data were retrieved from the GEO database and processed similarly to the ATA data processing procedure described in RNA-Chrom. The list of data is presented in Supplementary Tables 1 and 2. Only those RNAs that showed more than 1000 chromatin contacts in each replicate, which provides sufficient statistical power for peak detection by the BaRDIC program [18], were included in the analysis. Ribosomal RNAs were excluded from the analysis. For example, when this filter was applied in the experiments «RADICL, ES (NPM)» and «RADICL, ES (ActD)», less than 1000 RNAs and less than 50 percent of the contacts from the initial size of the selected replicates remained (Supplementary Fig. 1a, b). Given the significant overrepresentation of close contacts (RD-scaling), interactions located closer than 1 Mb from the genes encoding the corresponding RNAs were not included in further analysis.

### Using BaRDIC. Threshold selection

Like most genome-wide data, RNA-chromatin interactome data are characterized by high levels of non-specific signal («noise»). To identify significant interactions, specialized peak detection algorithms are used to identify statistically significant clusters of interactions at specific genomic loci.

The RNA-chromatin interactome data have a number of peculiarities. In particular, in all experiments, an increase in the density of contacts near the gene from which the RNA is transcribed is observed, with a gradual decrease in the signal with increasing distance from the parental gene along the chromosome. We call this phenomenon RD-scaling. On the other hand, the density of the observed contacts is affected by the degree of chromatin accessibility. To identify peaks in the data in this study, we use the BaRDIC algorithm [18], which we developed earlier, which takes into account these peculiarities. This algorithm uses a probabilistic estimate of the belonging of contacts in the chromatin locus to a peak or noise for each locus. Then, the Benjamini-Hochberg multiple testing correction (FDR) is applied, which controls the proportion of false positives (FP) based on the background distribution. However, in our case, a significant overlap of the signal and noise distributions leads to the loss of a significant proportion of true interactions when using a strict threshold on the FDR. To avoid this problem, we used a flexible selection criterion: for each RNA, we selected the top 10% of peaks with the lowest FDR. Since peak sizes could reach tens of kilobases due to data sparseness, all comparisons were performed at the level of individual contacts intersecting these peaks.

For the analysis of ATA data, BaRDIC was launched with default parameters.When processing OTA experiments with better contact coverage, the following parameters were set: --trans_min 400 pb, --cis_start 100 pb, --trans_step 50 pb. The background was calculated from the input data with conversion to BedGraph, the window size was 1000 bp.

### Chromatin potential

Almost all studies devoted to ATA experiments have noted that the number of RNA contacts with chromatin is linearly dependent on the expression level of the corresponding RNA. Normalization to the expression level allows us to isolate RNAs that demonstrate an increased tendency to interact with chromatin — those molecules whose frequency of contacts significantly exceeds that expected at a given expression level.

To assess the propensity of RNA to contact chromatin, we introduce the concept of *chromatin* potential. Let us consider all RNAs for which contacts are observed. Let *N*_*c*_ – the total number of contacts (library size) in the ATA experiment; *N*_*e*_ – the total number of reads in the RNA-seq experiment; 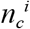 – the number of contacts of a particular i-th RNA in the ATA experiment, 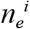 – the number of reads of a particular i-th RNA in the RNA-seq experiment. To compare these observations, we apply the Z-test of proportions. For each i-th RNA, we calculate the Z-statistic (1):

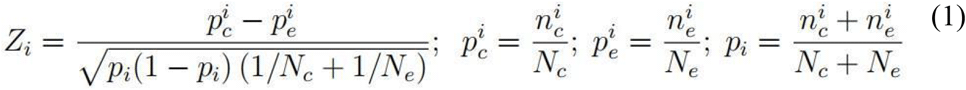

The Z statistic has a standard normal distribution and therefore can be used to estimate the p-value and the Benjamini-Hochberg FDR. The value of the Z statistic will be called the chromatin potential.

However, the following circumstances should be kept in mind. Firstly, such an analysis requires chain-oriented total RNA sequencing with rRNA depletion. Secondly, this analysis is applicable only to long RNAs, since standard RNA-seq data do not allow adequate assessment of the expression level of RNA shorter than 100 nucleotides [19].

## RESULTS

### Chromatin potential study

In the «Materials and methods» section we introduced the concept of chromatin potential. In this section we examine how the use of chromatin potential («chP») in the analysis of ATA data allows filtering out RNAs with predominantly random contacts with chromatin, as well as the dependence of chromatin potential on the number of contacts. Figure 2a shows the dependence of chromatin potential on the number of contacts for the RED-C experiments (other ATA experiments – Supplementary Figure 2).

**Figure 2.**
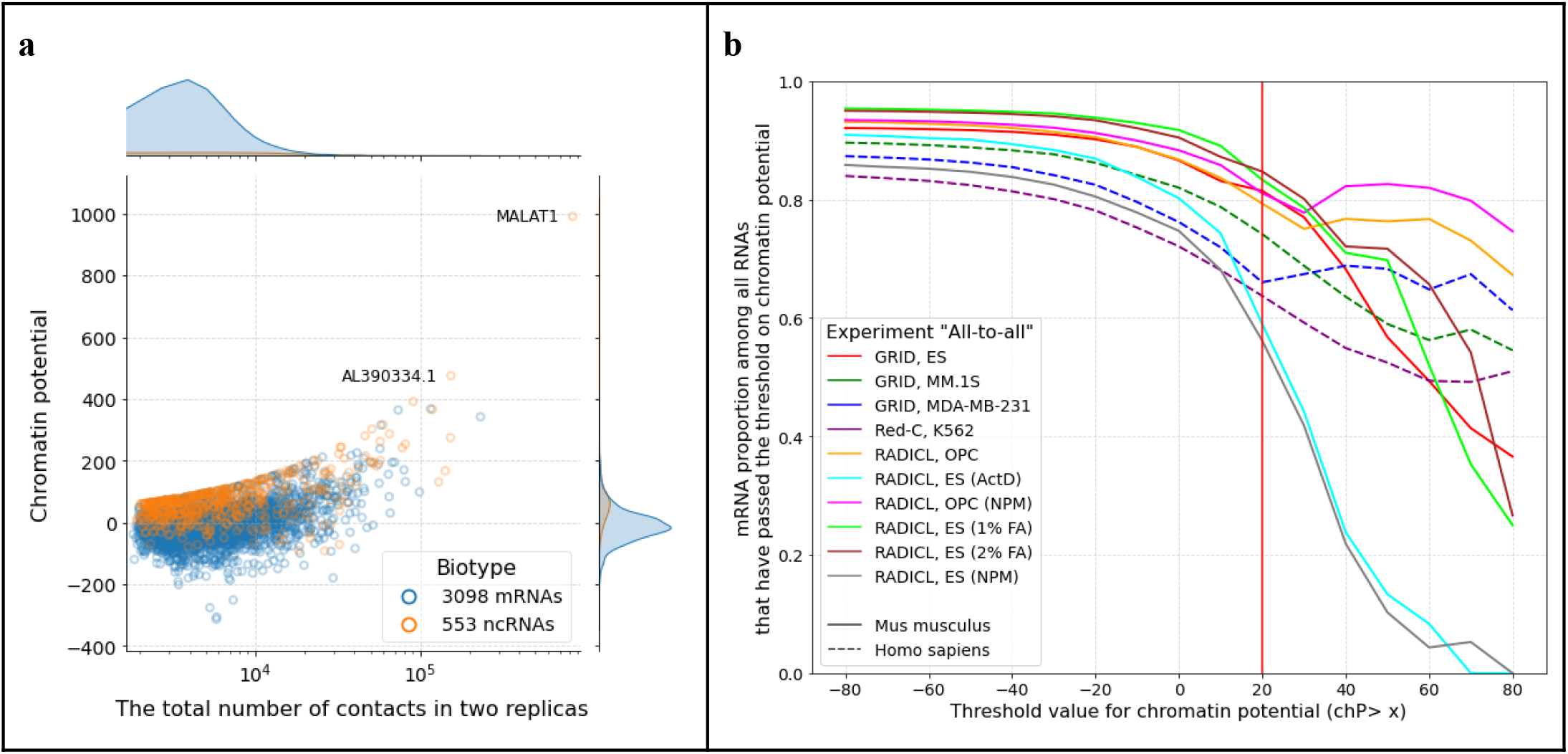
**a**: Chromatin potential as a function of RNA contact number in RED-C K562 experiment. Blue – protein-coding RNAs, orange – non-coding RNAs. **b**: mRNA fraction as a function of chromatin potential threshold (chP > x) for different experiments. ActD – actinomycin D treatment, NPM – proteinase K treatment, 1% FA – treatment with 1% formaldehyde cross-linking agent, 2% FA – treatment with 2% formaldehyde cross-linking agent.

In almost all genome-wide studies of RNA-chromatin interactions, a significant predominance of contacts of protein-coding RNAs (mRNAs) is observed. This is due to the fact that mRNAs, as a rule, have a higher expression level compared to ncRNAs. Chromatin potential should neutralize the effect of overrepresentation of contacts of highly expressed RNAs. If we assume that most mRNA contacts with chromatin are non-specific, then one would expect that non-coding RNAs should demonstrate a higher affinity for chromatin. Therefore, we looked at the dependence of the mRNA proportion on the chromatin potential threshold (Figure 2b). As expected, the graphs show a clear trend of a decrease in the mRNA proportion with an increase in the chromatin potential threshold. It is evident that at chromatin potential values of about 20, a sharp drop in the mRNA proportion is observed in almost all experiments. Supplementary Table 3 shows how the number of accepted RNAs depends on different chromatin potential thresholds.

Let us note some features. Firstly, with an increase in the threshold for chromatin potential, the proportion of protein-coding RNAs decreases (Figure 2b). Secondly, even at high thresholds for chromatin potential, quite a lot of protein-coding RNAs remain. This may be due to several circumstances, for example, many protein-coding genes contain functional non-coding RNAs in their intron regions (see, for example, [20, 21]). On the other hand, non-coding mRNA isoforms themselves can play a certain role in chromatin regulation [22]. In addition, it can be assumed that a significant proportion of the observed contacts are not specific. There is one more feature. In experiments on mouse cells, the drop in the proportion of mRNA is much more pronounced. Two RADICL experiments [13] on mouse embryonic stem cells stand out, where a much sharper drop in the proportion of mRNA is observed with an increase in the threshold for chP. These are experiments where cells were pre-incubated with actinomycin D, which suppresses transcription, or with proteinase K. Here it was assumed that only direct RNA-DNA interactions, not mediated by proteins, would remain. Apparently, these treatments of the material allow a significant number of non-specific contacts to be removed, and the use of chromatin potential allows the true contacts to be isolated.

### Comparison of replicas

Comparison of replicas. The next question we want to answer is how consistent are the replicas in the experiments. For each RNA, we use the following procedure. We split the genome into bins of size «*bin*» of nucleotides. We will call a bin concordant if for a given RNA this bin contains contacts from two sources (replicas or experiments), and discordant if it contains contacts in only one data set. Obviously, as the bin size increases, the proportion of concordant contacts will increase and will reach 1 at a bin size equal to the chromosome size. Therefore, we must also track the number of contacts in the bin. To estimate the randomness of coincidences, we can use the simplest model.

Let us assume that all RNA contacts with genomic DNA uniformly, then in one experiment the probability of at least one contact falling into a bin is estimated as 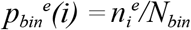, where *i* - the RNA number, *e* - the experiment (replica) number, 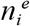 - the number of bins with contacts of the i-th RNA, *N*_*bin*_ – the total number of bins. Here we neglect data biases and assume that the bin size is small enough. To avoid the influence of RD-scaling, we select bins that are more than 1 Mb away from the i-th RNA gene. Then the probability that contacts from two experiments *a,b* will fall into one bin will be equal 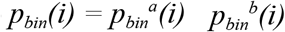. A rough estimate of the probability of observing *k* matching bins can be made using the Bernoulli distribution (2):

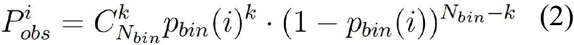

This allows us to make a probabilistic assessment of the correspondence of replicas or experiments. We define *λ(i) = (n*_*a*_*n*_*b*_*/N*_*bin*_). For λ ≥ 10 we can apply the normal approximation and estimate the probability of such an event, provided that the replicas have independent contacts (3):

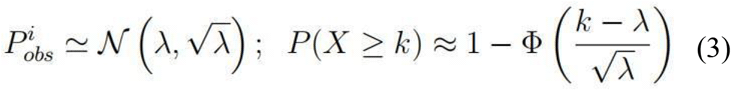

For λ < 10 we use the Poisson approximation (4):

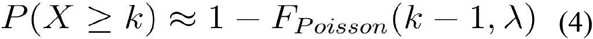

We tested the consistency of replicates for the ATA experiments. For this purpose, we selected RNAs that have more than 1000 contacts in each replicate, given a certain bin size (1000 or 5000 nucleotides). The RD-scaling filter was not applied to the RNA selection, but was applied to the contact selection. Table 1 shows the number of RNAs that have concordant bins between replicates with an FDR of less than 0.05. In this case, we took either all RNA contacts with chromatin or only contacts from BaRDIC peaks. When selecting contacts, a filter for the distance from the RNA source gene equal to 1 Mb was applied. Concordance and peaks are independent tests; however, it turns out that the selection of contacts from peaks corresponds to the selection of contacts by concordance. This table highlights the GRID experiments for which the filter for the significance of concordance does not reduce the number of selected RNAs. In this table, it can be seen that for GRID experiments, the filter for concordance significance does not reduce the number of RNAs sampled.

**Table 1.** Number of RNAs with concordant bins in replicates. N_concord is the number of RNAs whose concordance has an FDR of less than 0.05. Only RNAs with more than 1000 contacts in each replicate were selected.

| Experiment | Initial amount of RNA | N_concord (all contacts) |  | N_concord (contacts from peaks) |  |
| --- | --- | --- | --- | --- | --- |
|  |  | Bin 1000bp | Bin 5000bp | Bin 1000bp | Bin 5000bp |
| Red-C, K562, H.sap | 3866 | 1912 | 2904 | 3335 | 3816 |
| GRID, MM.1S, H.sap | 4184 | 4184 | 4184 | 4184 | 4184 |
| GRID, MDA_MB_231, H.sap | 5497 | 5497 | 5497 | 5497 | 5497 |
| GRID, ES, M.mus | 5142 | 5135 | 5133 | 5138 | 5141 |
| RADICL (2% FA), ES, M.mus | 2920 | 1916 | 2676 | 2357 | 2862 |
| RADICL, OPC, M.mus | 2777 | 2090 | 2659 | 2364 | 2746 |
| RADICL (ActD), ES, M.mus | 744 | 387 | 652 | 586 | 732 |
| RADICL (NPM), OPC, M.mus | 4009 | 544 | 1567 | 2264 | 3039 |
| RADICL (1% FA), ES,<br>M.mus | 2196 | 1599 | 2066 | 1913 | 2171 |
| RADICL (NPM), ES,<br>M.mus | 792 | 792 | 792 | 791 | 791 |

The analysis of the contact concordance gives us an idea of the completeness of the data. It is important to note that the complete data set includes both specific and non-specific interactions. Figure 3 and Supplementary Figure 3 show that in the ATA experiments, even with a bin size of 5000 bp, the concordance level is not very high and is 5-10%, and the concordance increases with the number of contacts. In addition, the concordance increases with the chromatin potential. From this we can conclude that the data are sufficiently complete only when the number of contacts exceeds 10 thousand, and the accuracy of contact determination on DNA is about 5 thousand nucleotides. Another feature to note is that some mRNAs have very concordant contacts, which can be interpreted as a high level of non-specificity, or that the mRNA has many specific contacts.

**Figure 3.**
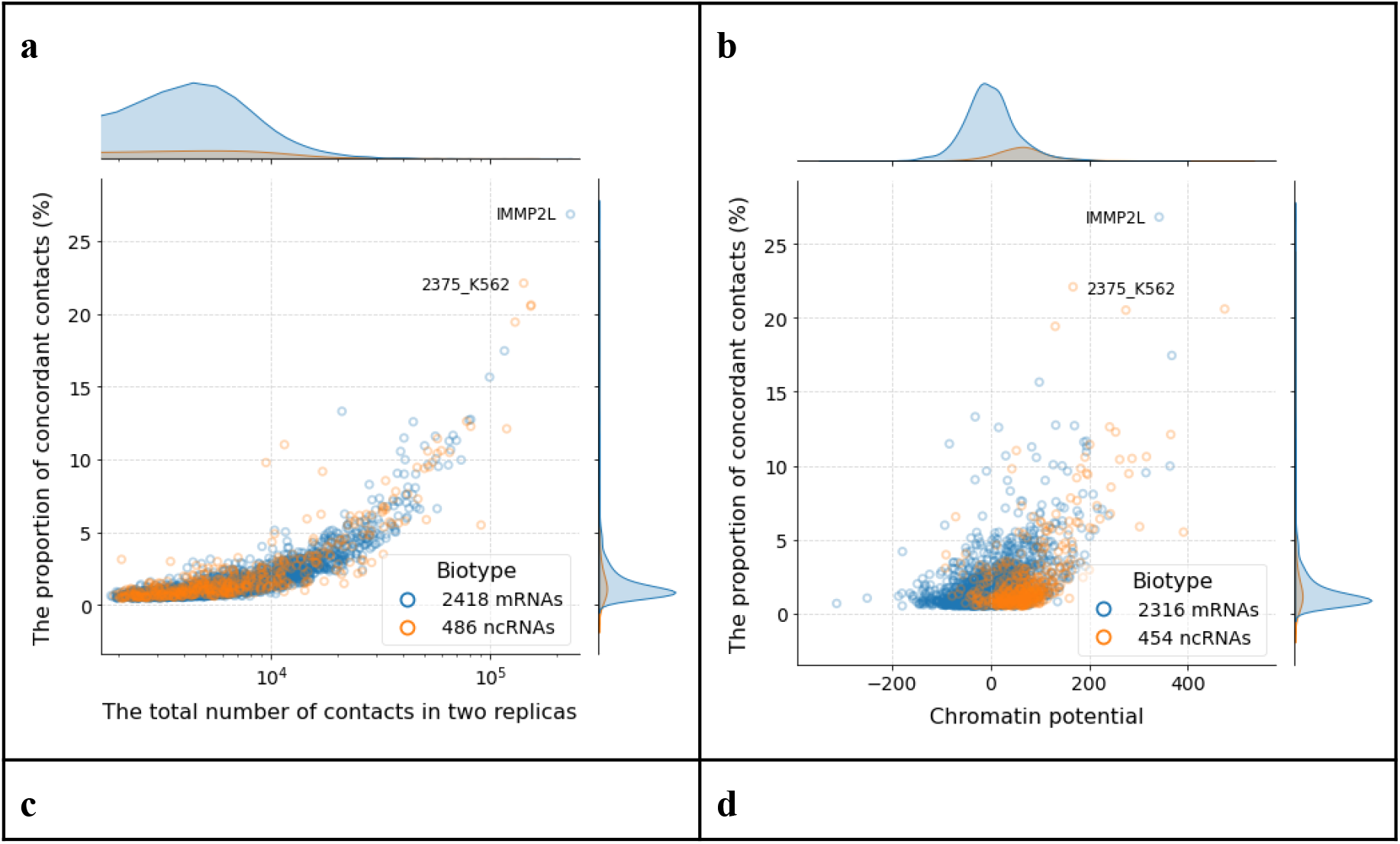
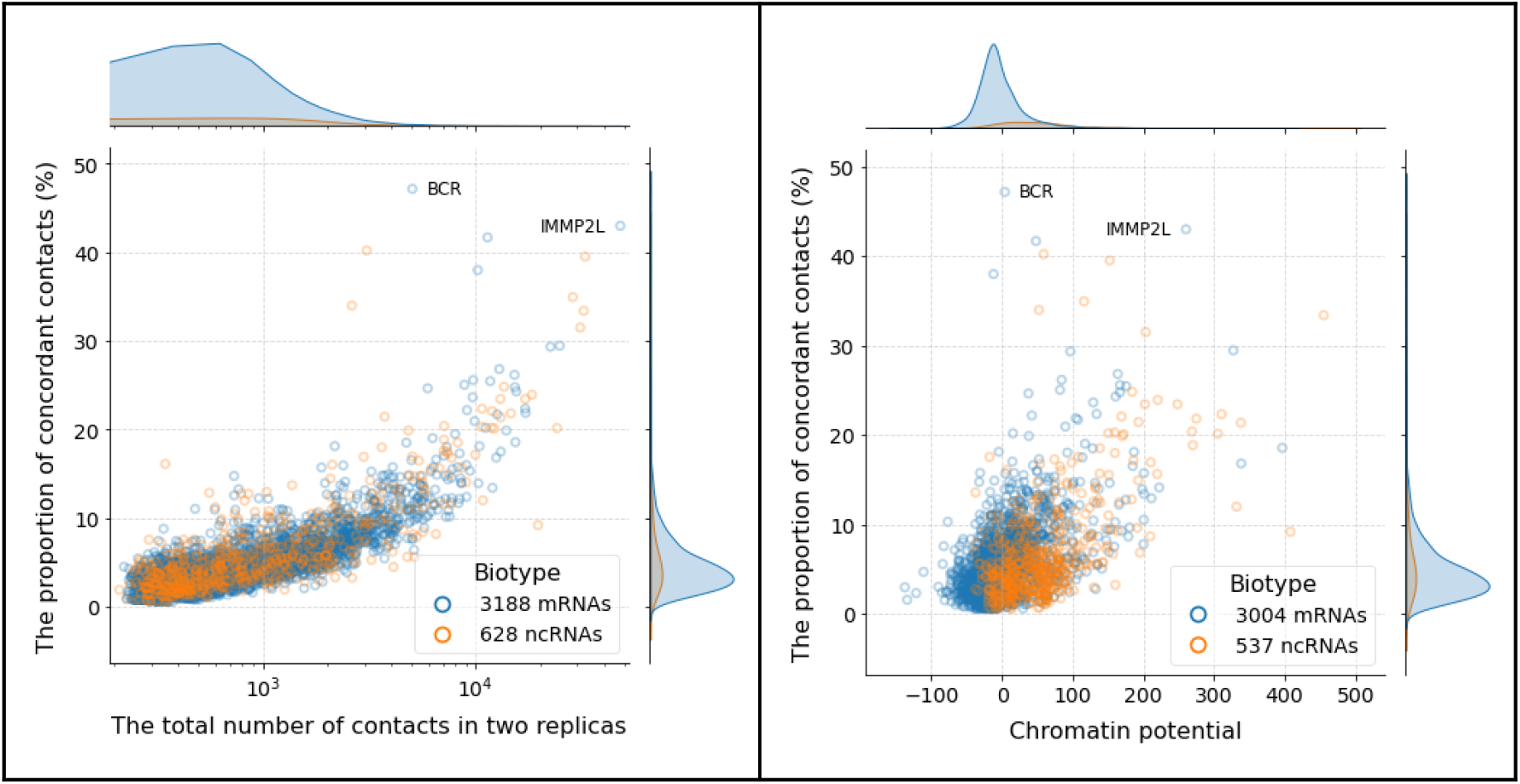
**a, b** presents the dependences of the concordance of replicas on the number of contacts and on the chromatin potential for a bin of 5000 nucleotides; **c, d** – the same, but for contacts from the peaks for the RED-C data on K-562 cells. MALAT1 is not shown on the graph, since this RNA has an extreme value of the chromatin potential and the proportion of concordant contacts: 991 and 58.2% - in figures a, b; 740 and 71.9% - in figures c, d.

In order to get an idea of the concordance level for different experiments, we calculated the median of the concordance level distribution (Supplementary Tables 4 and 5). It should be noted that the median concordance of replicates in the GRID data is high, sometimes reaching 39% (80% at the peaks), while in the Red-C and RADICL data the median concordance rarely exceeds 10%. For the GRID experiments, the processed data are publicly available. Apparently, such a high percentage of concordance is explained by the features of the experimental protocol and the primary data processing. Another observation is that the concordance rate of protein-coding RNAs does not differ from that of ncRNAs.

Next, we estimated the concordance level of replicates in experiments with individual RNAs (OTA) (Table 2). There are two peculiarities here. First, in most experiments the concordance level is over 90%, which means good data completeness. However, if we use only contacts from peaks, the concordance level drops by almost half. The peak extraction program looks for contact clusters and discards sporadic single contacts, which are apparently non-specific. Therefore, this drop in the concordance level is due to a fairly high proportion (more than half) of non-specific contacts.

**Table 2.** The proportion of concordant contacts in OTA replicates (%). d0, d3 and d7 - 0, 3 and 7 days of cell differentiation, respectively. p-value < 0.05.

| RNA | Experiment | Bin=1000bp |  | Bin=5000bp |  |
| --- | --- | --- | --- | --- | --- |
|  |  | All contacts | Contacts from peaks | All contacts | Contacts from peaks |
| JPX | CHART, ES d0 (GSM4278791, GSM4278795) | 99,5 | 53,3 | 100,0 | 78,2 |
| JPX | CHART, ES d3 (GSM4278799, GSM4278803) | 99,3 | 36,8 | 100,0 | 70,6 |
| JPX | CHART, ES d7 (GSM4278807, GSM4278811) | 99,2 | 44,5 | 100,0 | 75,0 |
| MAL AT1 | ChIRP, ES, genotype: Ythdc1-cKO (conditional); treatment: DMSO, (GSM4669091, GSM4669092) | 79,9 | 26,9 | 99,6 | 50,6 |
| MAL AT1 | ChIRP, ES, genotype: Mettl3-WT, (GSM4875651, GSM4875652) | 92,4 | 40,9 | 99,9 | 66,3 |

Our analysis of the replica concordance in RNA-chromatin interactome experiments showed that the ATA-type experiments generally have low concordance rates of less than 10%. This leads to the conclusion that the data are substantially incomplete and a significant proportion of contacts are missed. On the other hand, the concordance rates in OTA experiments are generally greater than 90%, allowing these experiments to be used as a gold standard.

### Comparison of experiments

The concordance analysis of replicates of the single RNA interactome (OTA) data showed their high consistency, so we use these data to evaluate the data of the same RNAs in genome-wide assays (ATA). Unfortunately, such an analysis is very limited, since we need data obtained for the same cell lines and with a sufficiently large number of contacts in the ATA data. In the comparative analysis of experiments, we used bins of 5000 nucleotides. Since the number of contacts for a given RNA differs by several orders of magnitude, we will consider the proportion of concordant contacts of the corresponding RNA in the ATA data as a measure of concordance - the number of concordant contacts in the ATA experiment divided by the total number of contacts of the corresponding RNA in the ATA data. In this case, we will take into account both all contacts and contacts from the BaRDIC peaks. To increase the completeness of the data, we combined sets of contacts in replicates in this analysis. The results for the ncRNAs MALAT1 and JPX are presented in Tables 3 and 4.

**Table 3.** Percentage of concordant MALAT1 RNA contacts with chromatin in ATA data compared with MALAT1 RNA contacts from OTA experiments (contacts from BaRDIC peaks) in mouse embryonic stem cells. The result for ATA contacts from BaRDIC peaks is presented in brackets. Bin size is 5000bp, p-value < 0.05, d0, d3 and d7 - 0, 3 and 7 days of cell differentiation, respectively.

|  | Number of contacts in ATA data | RAP |  | ChIRP |  |  |
| --- | --- | --- | --- | --- | --- | --- |
|  |  | pSM33 ES, DMSO 1 hour | V6.5 ES | ES, Ythdc1-cKO; DMSO | ES, Mettl3-WT | E14 ES |
| GRID, ES, M.mus | 522 741<br>(109 371) | 38,0 (46,5) | 55,8<br>(61,6) | 50,6<br>(51,0) | 58,8<br>(58,3) | 53,8<br>(57,0) |
| RADICL (1% FA), ES, M.mus | 636 802<br>(138 422) | 42,9 (56,2) | 61,5<br>(74,1) | 50,4<br>(49,6) | 58,9<br>(57,2) | 55,1<br>(58,6) |
| RADICL (2% FA), ES, M.mus | 484 878<br>(99 985) | 41,5 (51,0) | 59,7<br>(69,0) | 50,7<br>(49,6) | 59,2<br>(58,2) | 54,9<br>(56,8) |

**Table 4.** Percentage of concordant JPX RNA contacts with chromatin in ATA data (all contacts) compared with JPX RNA contacts from OTA experiments (contacts from BaRDIC peaks) in mouse embryonic stem cells. Concordance p-value is given in brackets. Bin size is 5000bp, d0, d3 and d7 – 0, 3 and 7 days of cell differentiation, respectively.”

|  | Number of Jpx contacts in ATA data | CHART, ES |  |  |
| --- | --- | --- | --- | --- |
|  |  | d0 | d3 | d7 |
| GRID, ES, M.mus | 459 | 57,1 (0,05) | 61,9 (0,22) | 63,6 (0,002) |
| RADICL (1% FA), ES, M.mus | 341 | 57,2 (0,09) | 62,8 (0,15) | 61,0 (0,03) |
| RADICL (2% FA), ES, M.mus | 332 | 54,5 (0,24) | 56,6 (0,65) | 63,9 (0,005) |

When analyzing these comparisons, it should be borne in mind that although the data were obtained for the same or similar cell lines, the conditions of their cultivation differed in different OTA experiments. Nevertheless, we can draw the following conclusions. First, for the MALAT1 data, the proportion of contacts in the ATA data that coincide with the OTA data is quite high and amounts to 50%. This is primarily due to the fact that for MALAT1 ncRNA, the number of observed contacts in ATA is very large. As noted earlier, for replicates in ATA experiments, the concordance of MALAT1 ncRNA with MALAT1 ncRNA is also quite high. This value can serve as an estimate of the proportion of specific contacts. On the other hand, the remaining 50% should be attributed to nonspecific contacts.

For the JPX ncRNA the situation is different – the number of contacts of this RNA in the ATA data is small and this RNA was not involved in our previous analysis. The concordance of contacts with the OTA data is only about 60%. We can therefore estimate the proportion of non-specific contacts as 40%. It is known that the JPX ncRNA is associated with the XIST ncRNA [23], as expected, the main proportion of contacts of this RNA is localized on the X chromosome and the density of contacts of this RNA on the X chromosome is quite high. Using peaks would significantly increase the specificity estimate, but the number of contacts of the JPX RNA is insufficient to trigger BaRDIC.

To further assess the specificity, we compared data from OTA experiments for different ncRNAs in human (Figure 4) and mouse (Supplementary Figure 4) cells. As a measure of the similarity of interaction maps, we assessed the ratio of concordant contacts to the total number of identified interactions. Several bright triangles corresponding to certain RNAs stand out in the heat map: MALAT1, NEAT1, TERC, and others. The upper part of the map as a whole shows mutual concordance of contacts for many ncRNAs.Against this background, there are quite bright dots showing concordance of contacts of different RNAs with chromatin. We noticed a significant concordance between the contact profiles of MALAT1, NEAT1, and SRA1. Long ncRNAs MALAT1/NEAT1 are predominantly localized near actively transcribed genes, while SRA1 RNA functions as a co-activator of steroid receptors. TERC contacts are also concordant with the poorly characterized ncRNA LINC00520. The functional relationship of this RNA with TERC is unknown, which may indicate low specificity of the data. If we accept that most RNA interactions with chromatin are mediated by proteins, then the low specificity of these contacts can be explained not by the peculiarities of the experimental methods, but by the relatively low specificity of the RNA-binding domains of proteins (see, for example, [24, 25]).

**Figure 4.**
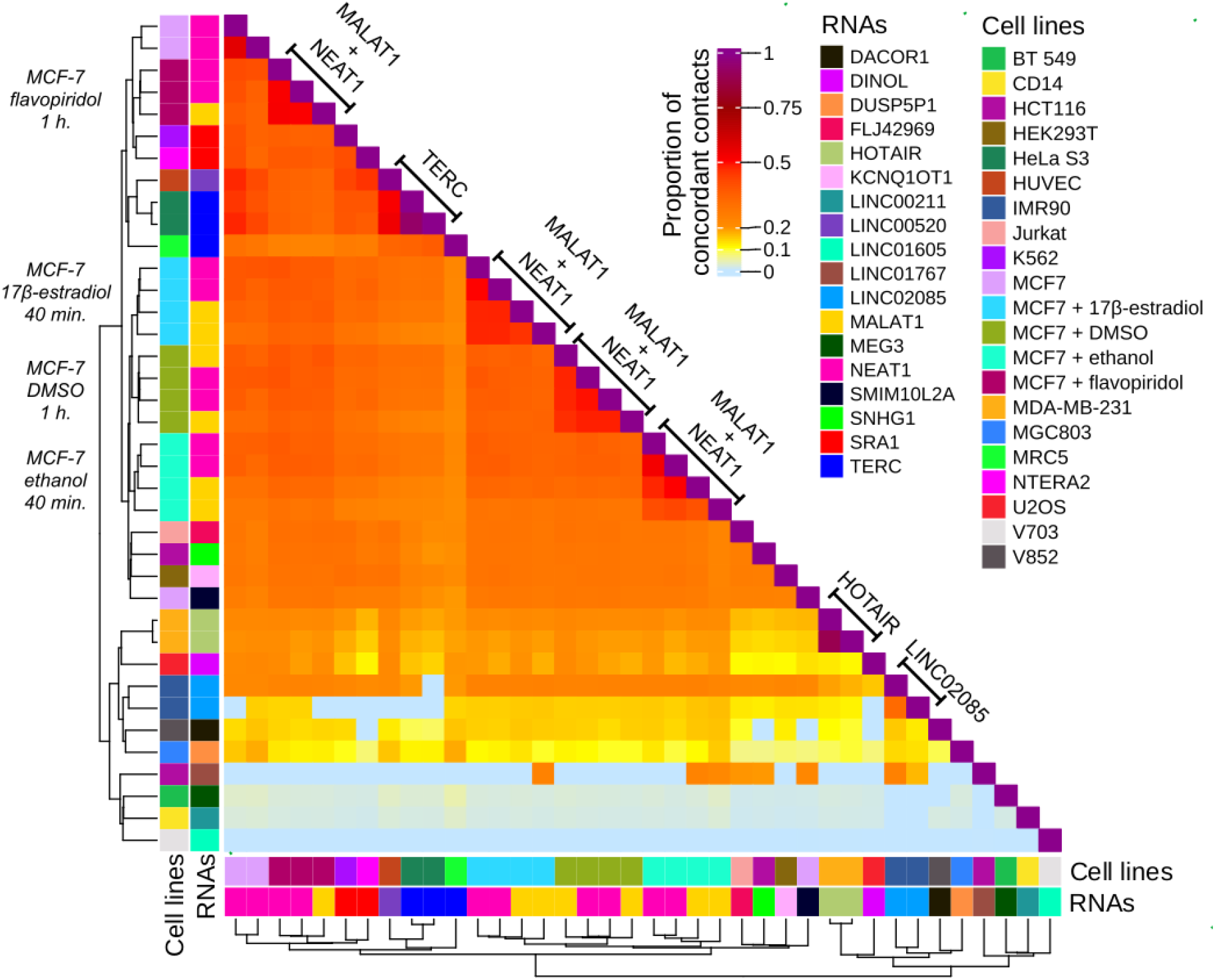
Heatmap showing the proportion of concordant contacts (from BaRDIC peaks) from «one-to-all» experiments for human cell lines. Non-significant enrichments (p-value > 0.05) are set to zero. Clustering is performed by cell types and RNAs used in the experiment. The bin size is 1000 bp.

The OTA data (Supplementary Figure 4) for mouse are mostly devoted to XIST. Bright spots from different RNAs are usually associated with X chromosome inactivation. For example, JPX RNA is an activator of XIST [23] and their association is not surprising.

## CONCLUSION

In this work, we performed a comparative analysis of RNA chromatin interactome data. We compared genome-wide RNA chromatin interactome (ATA) data with RNA-seq data and introduced the concept of chromatin potential, a numerical characteristic of individual RNAs that shows the extent to which the number of contacts of a given RNA exceeds that expected from RNA sequencing data. It was shown that setting a threshold for this value allows for a significant reduction in the proportion of protein-coding RNAs in the interactome, thereby reducing the number of nonspecific contacts.

Next, we examined the correspondence of contact maps between replicates and introduced a numerical characteristic of replicate concordance. We assume that this measure characterizes the completeness of the data. Statistical analysis showed that the concordance requirement has virtually no effect on RNA selection: virtually all RNAs with more than 1,000 contacts are statistically significantly concordant. It turned out that for full-genome data, concordance varies widely from 1% to 30%, indicating high variability of these data. At the same time, both mRNA and ncRNA have a similar concordance level.

The OTA experiments show very high replica concordance (around 90%), with concordance decreasing significantly at peaks but remaining quite high. This allowed us to use these data as a gold standard. Comparison of OTA data with ATA data allows us to estimate the specificity of ATA data.

## Accepted abbreviations

OTA («one-to-all»): experimental methods that allow determining contacts of previously known RNA with chromatin;
ATA («all-to-all»): experimental methods aimed at determining all possible RNA-DNA contacts in a cell.

