## Supplementary Tables for "COMPARATIVE ANALYSIS OF THE RNA-CHROMATIN INTERACTOME DATA": Supplementary_materials.pdf

**Supplementary Table 1.** ATA data. ActD - treatment with actinomycin D, NPM - treatment with proteinase K, 1% FA - treatment with the cross-linking agent formaldehyde at a concentration of 1%, 2% FA - treatment with the cross-linking agent formaldehyde at a concentration of 2%.

| **ATA data (experimental method, cell line, organism)** | **Replica numbers** | **Number of contacts** |
|---|---|---|
| GRID, MM.1S, Homo sapiens | GSM2188868, GSM2188869 | 37 817 797 |
| Red-C, K562, Homo sapiens | GSM4041591, GSM4041595 | 41 777 653 |
| RADICL, OPC, Mus musculus | GSM3852782-GSM3852783, GSM3852784-GSM3852785 | 27 898 478 |
| RADICL, ES (ActD), Mus musculus | GSM3852772-GSM3852773, GSM3852774-GSM3852775 | 13 586 144 |
| RADICL, OPC (NPM), Mus musculus | GSM3852788-GSM3852789, GSM3852790-GSM3852791 | 42 648 037 |
| RADICL, ES (1% FA), Mus musculus | GSM3852760-GSM3852761, GSM3852762-GSM3852763 | 28 609 319 |
| RADICL, ES (2% FA), Mus musculus | GSM3852766-GSM3852767, GSM3852768-GSM3852769 | 28 849 211 |
| RADICL, ES (NPM), Mus musculus | GSM3852776-GSM3852777, GSM3852778-GSM3852779 | 10 444 053 |
| GRID, ES, Mus musculus | GSM2396700, GSM2396701 | 59 958 702 |
| GRID, MDA_MB_231, Homo sapiens | GSM2188866, GSM2188867 | 63 227 196 |

**Supplementary Table 2.** RNA-seq data with ribosomal RNA depletion.

| **Cell line, organism** | **Replica numbers** |
|---|---|
| MDA-MB-231, Homo sapiens | GSM7143069-GSM7143071 |
| MM.1S, Homo sapiens | GSM5788444 (SRR17510863, SRR17510864), GSM5788446 (SRR17510859, SRR17510860) |
| H1 ES, Homo sapiens | GSM3630264, GSM3630265 |
| K562, Homo sapiens | GSM4744788, GSM4744789, GSM4744790 |
| OPC, Mus musculus | GSM3034716, GSM3034717, GSM3034718 |
| ES, Mus musculus | GSM4775002, GSM4775004 |

**Supplementary Table 3**. The number of RNAs remaining when using a chromatin potential (chP) filter. ActD - actinomycin D treatment, NPM - proteinase K treatment, 1% FA - treatment with 1% formaldehyde crosslinking agent, 2% FA - treatment with 2% formaldehyde crosslinking agent.

| Experiment | chrP > 0 | | chrP > 20 | | chrP > 50 | |
|---|---|---|---|---|---|---|
| | mRNAs | ncRNAs | mRNAs | ncRNAs | mRNAs | ncRNAs |
| Red-C, K562, H.sap | 1256 | 486 | 769 | 438 | 362 | 328 |
| GRID, MM.1S, H.sap | 1705 | 374 | 1002 | 349 | 338 | 235 |
| GRID, MDA_MB_231, H.sap | 1806 | 564 | 890 | 458 | 250 | 116 |
| GRID, ES, M.mus | 2201 | 340 | 691 | 157 | 63 | 48 |
| RADICL, OPC, M.mus | 1056 | 161 | 564 | 147 | 129 | 40 |
| RADICL, ES (ActD), M.mus | 219 | 54 | 56 | 39 | 2 | 13 |
| RADICL, OPC (NPM), M.mus | 1658 | 219 | 861 | 201 | 195 | 41 |
| RADICL, ES (1% FA), M.mus | 891 | 80 | 285 | 57 | 30 | 13 |
| RADICL, ES (2% FA), M.mus | 1150 | 121 | 361 | 65 | 38 | 15 |
| RADICL, ES (NPM), M.mus | 263 | 89 | 105 | 82 | 4 | 35 |

**Supplementary Table 4**. Median percentage of concordant contacts of mRNA and ncRNA in different ATA experiments (bin size 5000 bp, RD-scaling filter 1 Mb). ActD - treatment with actinomycin D, NPM - treatment with proteinase K, 1% FA - treatment with cross-linking agent formaldehyde at a concentration of 1%, 2% FA - treatment with cross-linking agent formaldehyde at a concentration of 2%.

| Experiment | All mRNAs | mRNAs (chP > 20) | mRNAs (chP > 50) | All ncRNAs | ncRNAs (chP > 20) | ncRNAs (chP > 50) |
|---|---|---|---|---|---|---|
| GRID, ES, M.mus | 15.7 | 22.2 | 34.6 | 14.2 | 14.9 | 9.8 |
| GRID, MM.1S, H.sap | 10.4 | 11.9 | 14.3 | 10.1 | 10.3 | 10.6 |
| GRID, MDA_MB_231, H.sap | 28.5 | 32.4 | 38.6 | 28.6 | 28.9 | 31.6 |
| Red-C, K562, H.sap | 1.1 | 1.8 | 2.5 | 1.3 | 1.4 | 1.6 |
| RADICL, OPC, M.mus | 1.8 | 4.0 | 8.6 | 1.7 | 1.8 | 6.7 |
| RADICL (ActD), ES, M.mus | 1.1 | 3.1 | 0.0 | 1.4 | 2.5 | 11.4 |
| RADICL (NPM), OPC, M.mus | 1.0 | 1.5 | 2.8 | 1.0 | 1.0 | 2.0 |
| RADICL (1% FA), ES, M.mus | 1.2 | 3.4 | 11.3 | 1.1 | 2.9 | 7.0 |
| RADICL (2% FA), ES, M.mus | 1.6 | 4.3 | 13.5 | 1.6 | 2.4 | 9.0 |
| RADICL (NPM), ES, M.mus | 0.5 | 1.1 | 3.5 | 0.6 | 1.1 | 6.8 |

**Supplementary Table 5**. Median percentage of concordant contacts of mRNA and ncRNA in different ATA experiments (bin size 5000 bp, RD-scaling filter 1 Mb). Contacts from peaks. ActD - treatment with actinomycin D, NPM - treatment with proteinase K, 1% FA - treatment with cross-linking agent formaldehyde at a concentration of 1%, 2% FA - treatment with cross-linking agent formaldehyde at a concentration of 2%.

| Experiment | All mRNAs | mRNAs (chP > 20) | mRNAs (chP > 50) | All ncRNAs | ncRNAs (chP > 20) | ncRNAs (chP > 50) |
|---|---|---|---|---|---|---|
| GRID, ES, M.mus | 58.6 | 57.3 | 65.1 | 56.7 | 45.1 | 20.5 |
| GRID, MM.1S, H.sap | 46.9 | 44.5 | 44.6 | 44.6 | 43.1 | 39.7 |
| GRID, MDA_MB_231, H.sap | 81.6 | 73.8 | 69.8 | 81.1 | 77.4 | 64.7 |
| Red-C, K562, H.sap | 4 | 7.7 | 9.2 | 4.6 | 5.7 | 6 |
| RADICL, OPC, M.mus | 6 | 13.8 | 21.1 | 5.2 | 5.7 | 12.4 |
| RADICL (ActD), ES, M.mus | 4.2 | 10.2 | 10.3 | 5 | 7.6 | 19.6 |
| RADICL (NPM), OPC, M.mus | 3.1 | 5.5 | 8.2 | 3.1 | 3.4 | 6.5 |
| RADICL (1% FA), ES, M.mus | 3.8 | 9 | 22.4 | 3.2 | 5.5 | 10.2 |
| RADICL (2% FA), ES, M.mus | 5.6 | 14.6 | 29.5 | 4 | 4.9 | 12.5 |
| RADICL (NPM), ES, M.mus | 2.5 | 3.5 | 7.4 | 2.6 | 3.5 | 10.8 |

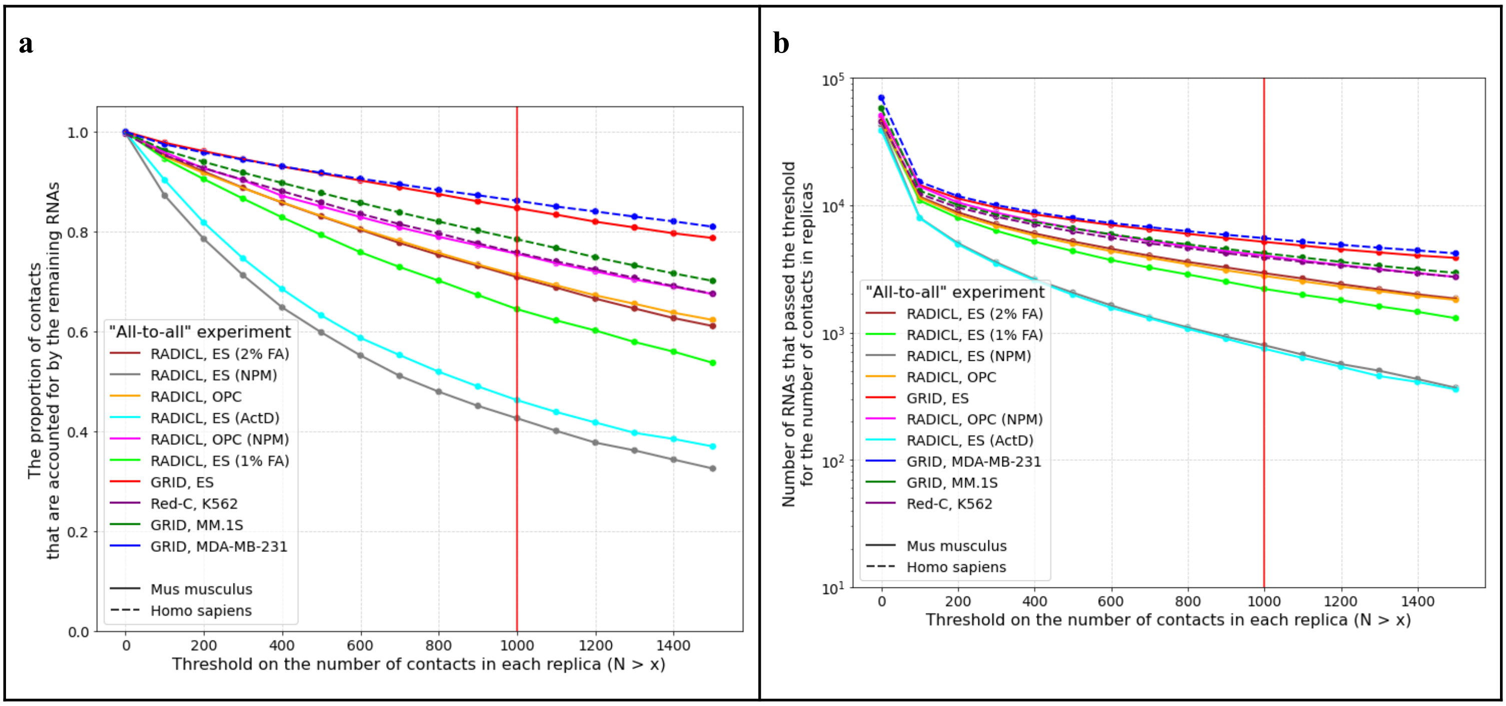


**Supplementary Figure 1. (a)** Contact fraction and **(b)** RNA amount as a function of contact threshold in each replicate for different experiments. ActD - actinomycin D treatment, NPM - proteinase K treatment, 1% FA - treatment with 1% formaldehyde cross-linking agent, 2% FA - treatment with 2% formaldehyde cross-linking agent.

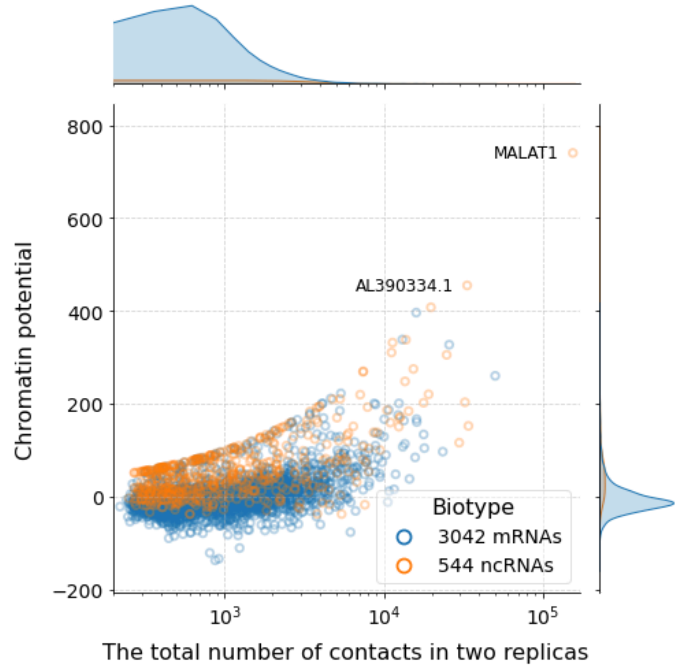


**A.** Red-C, K562, H.sap, contacts from peaks

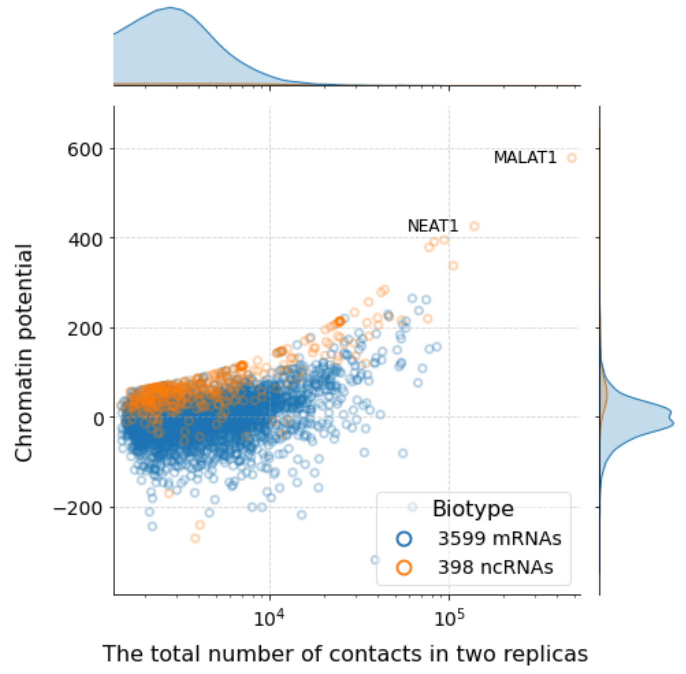


**B.** GRID, MM.1S, H.sap, all contacts

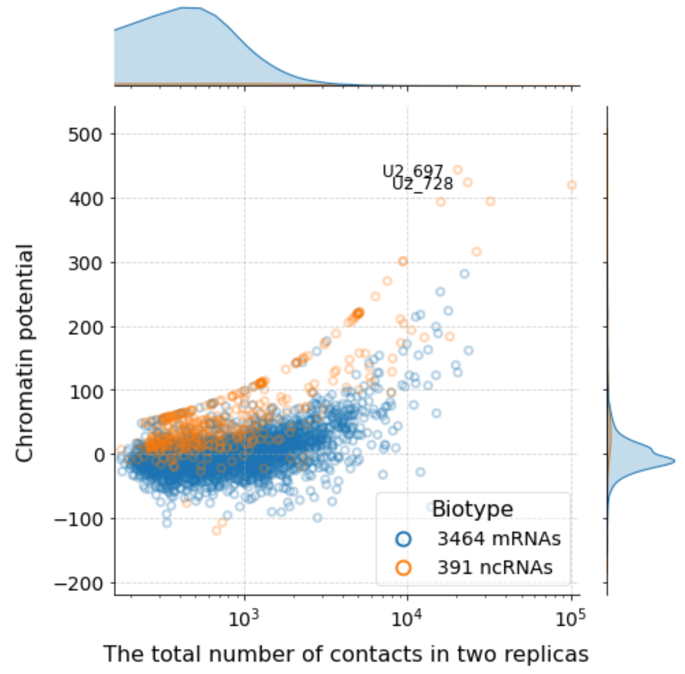


**C.** GRID, MM.1S, H.sap, contacts from peaks

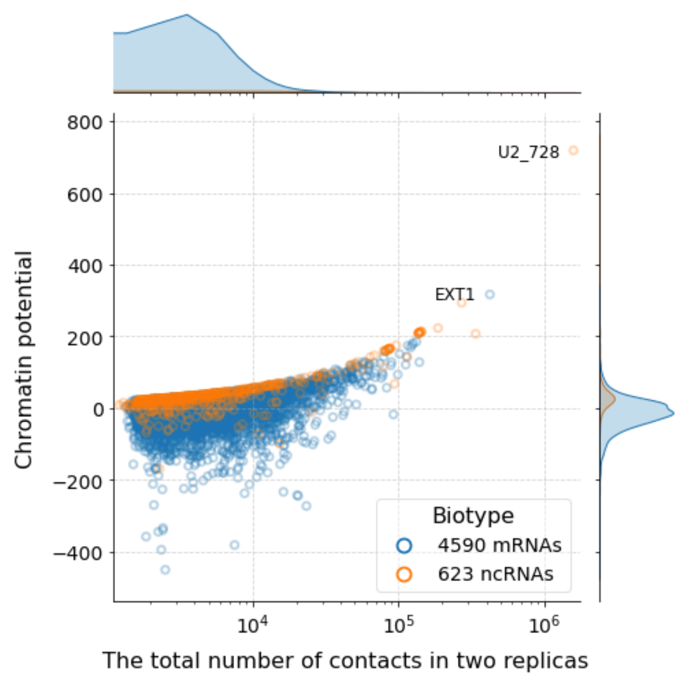


**D.** GRID, MDA_MB_231, H.sap, all contacts

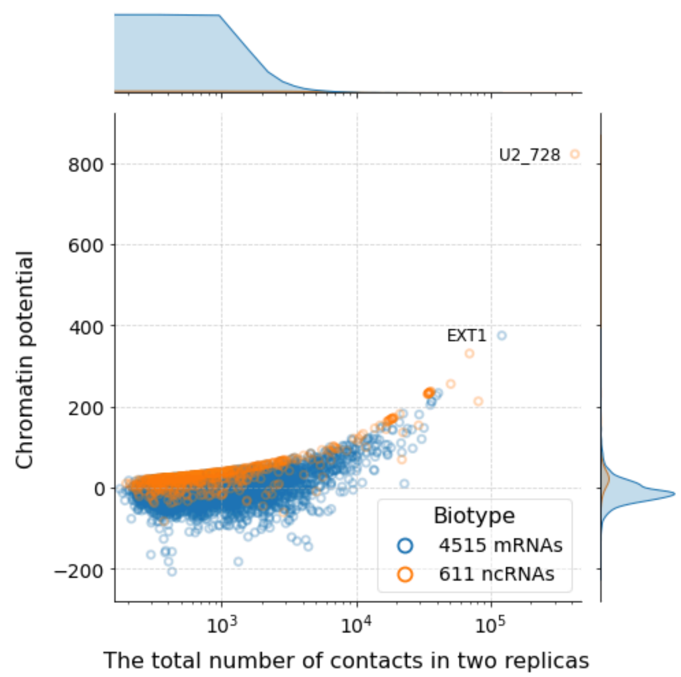


**E.** GRID, MDA_MB_231, H.sap, contacts from peaks

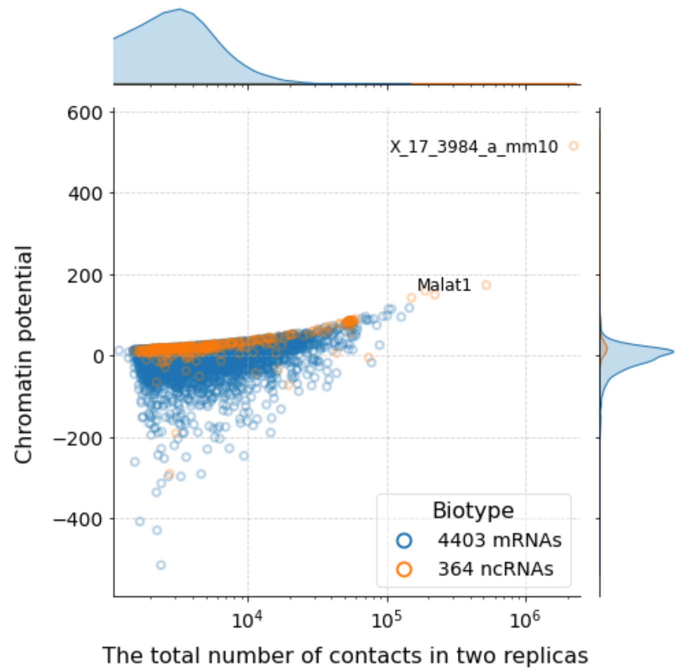


**F.** GRID, ES, M.mus, all contacts

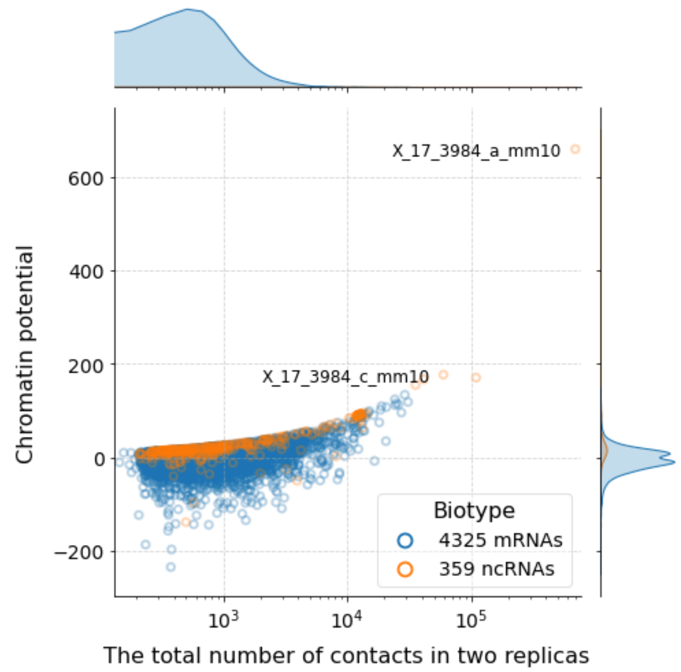


**G.** GRID, ES, M.mus, contacts from peaks

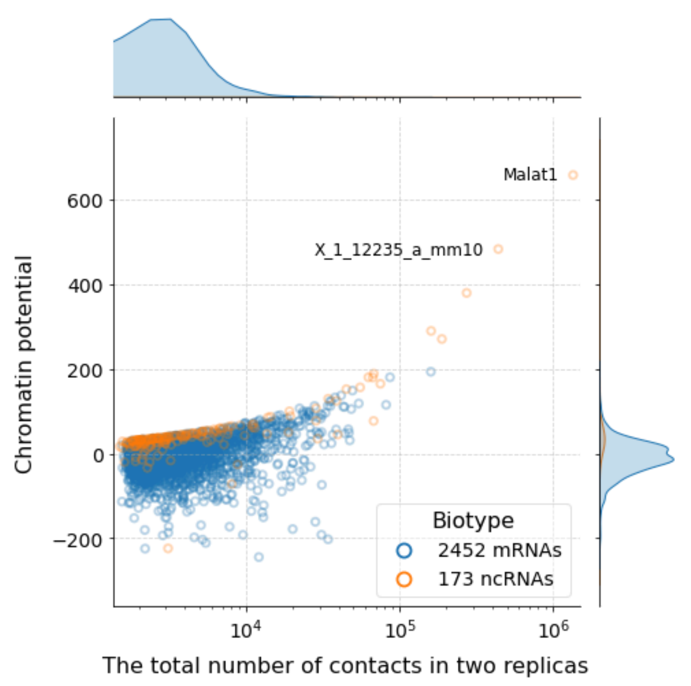


**H.** RADICL, OPC, M.mus, all contacts

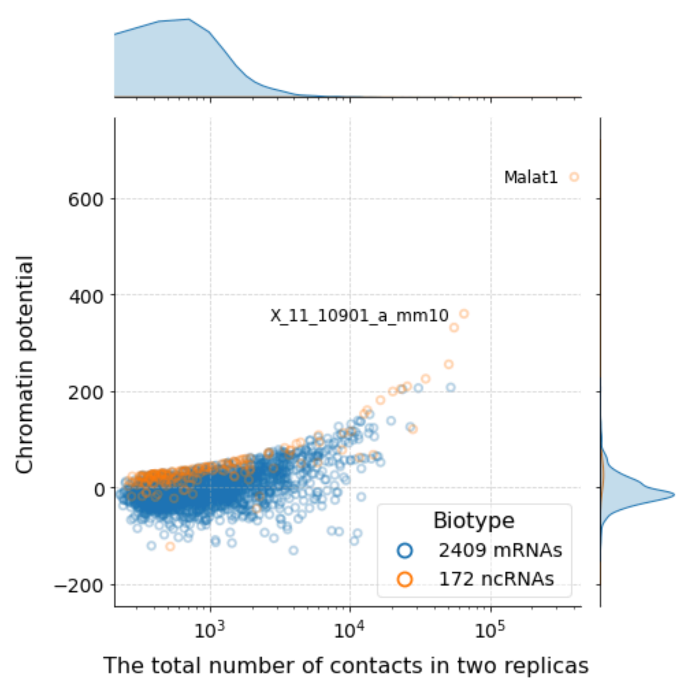


**I.** RADICL, OPC, M.mus, contacts from peaks

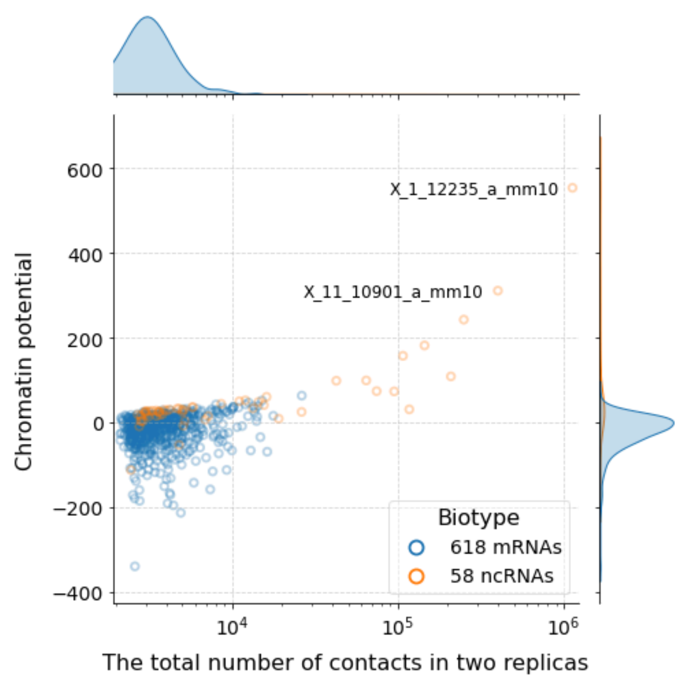


**J.** RADICL (ActD), ES, M.mus, all contacts

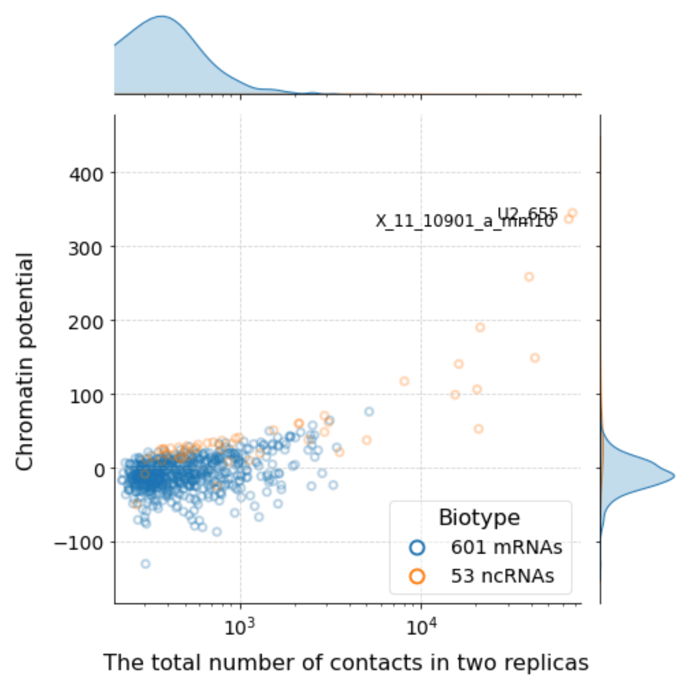


**K.** RADICL (ActD), ES, M.mus, contacts from peaks

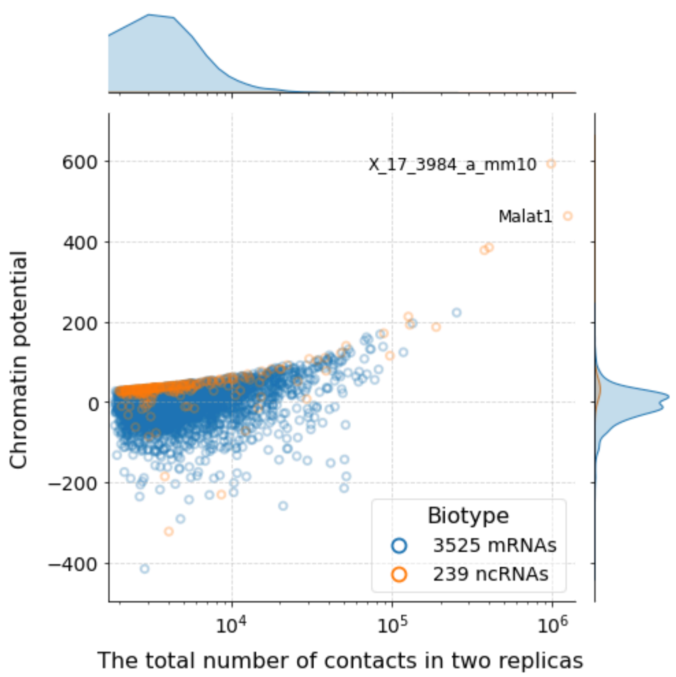


**L.** RADICL (NPM), OPC, M.mus, all contacts

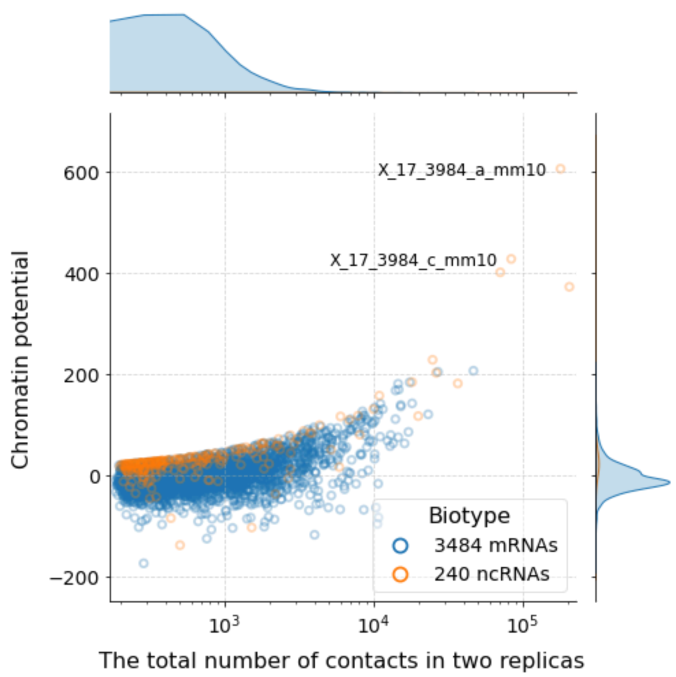


**M.** RADICL (NPM), OPC, M.mus, contacts from peaks

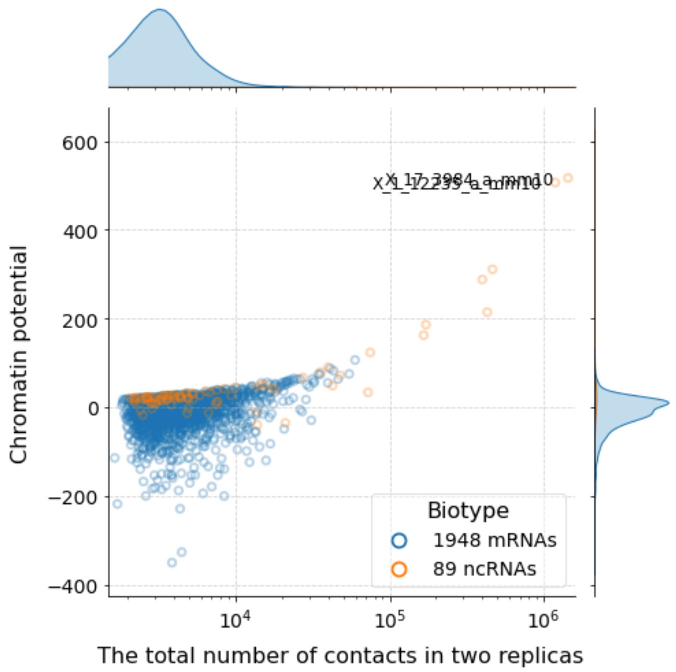


**N.** RADICL (1% FA), ES, M.mus, all contacts

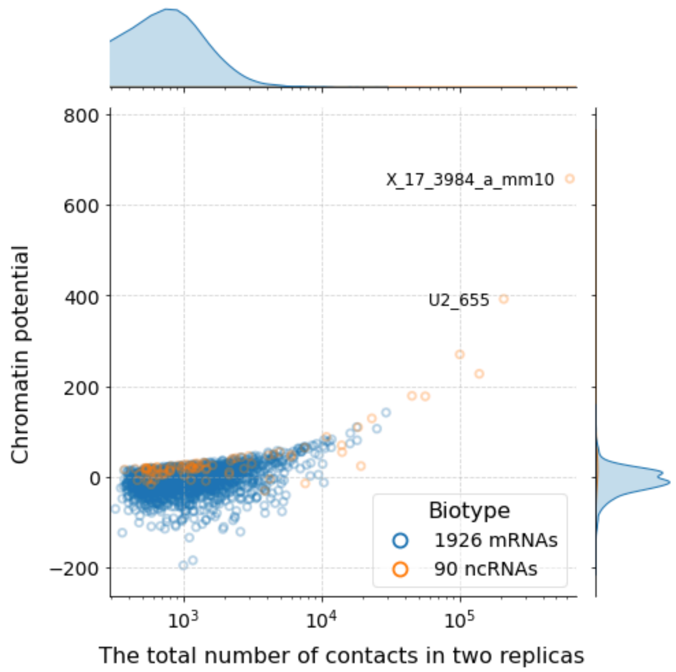


**O.** RADICL (1% FA), ES, M.mus, contacts from peaks

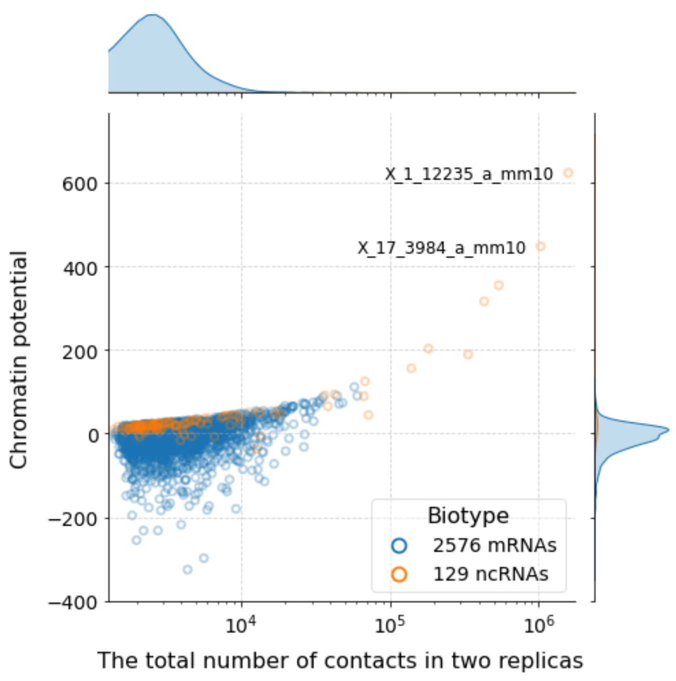


**P.** RADICL (2% FA), ES, M.mus, all contacts

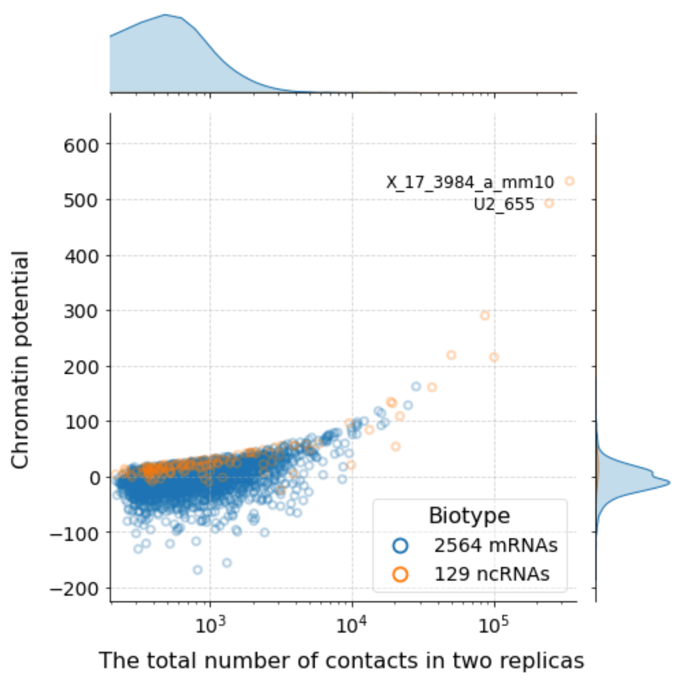


**Q.** RADICL (2% FA), ES, M.mus, contacts from peaks

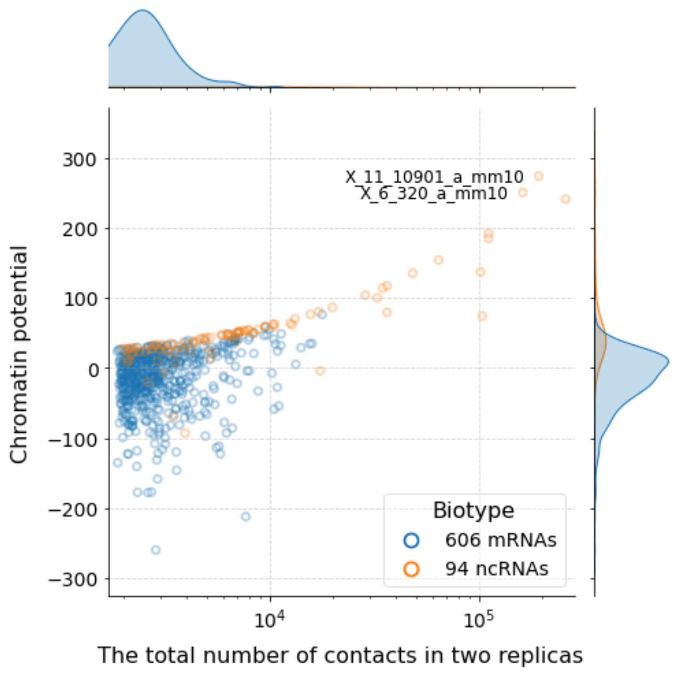


**R.** RADICL (NPM), ES, M.mus, all contacts

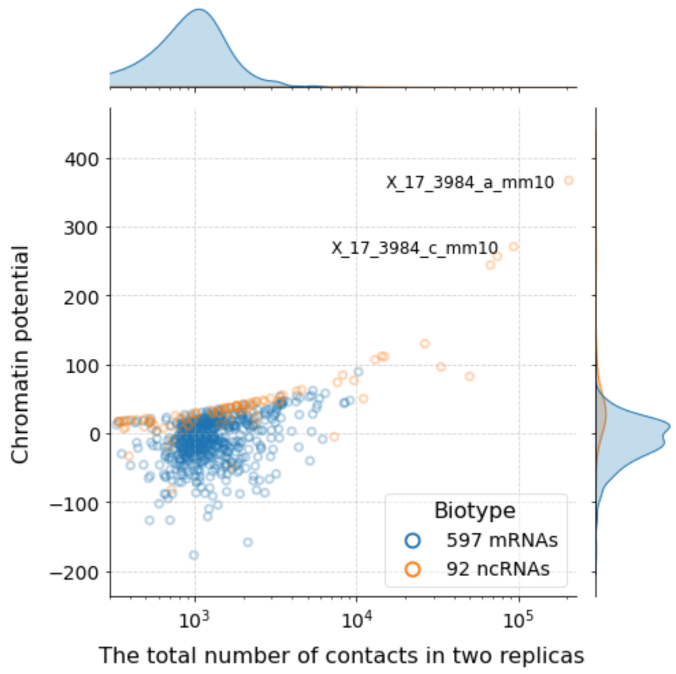


**S.** RADICL (NPM), ES, M.mus, contacts from peaks

**Supplementary Figure 2.** Dependence of chromatin potential on the number of contacts for all contacts and for contacts from peaks.

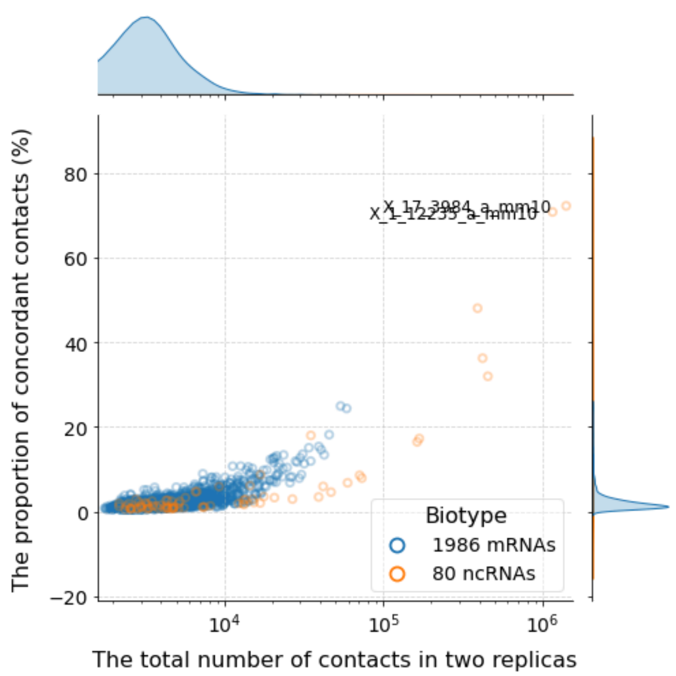


**A1.** RADICL (1% FA), ES, M.mus, all contacts

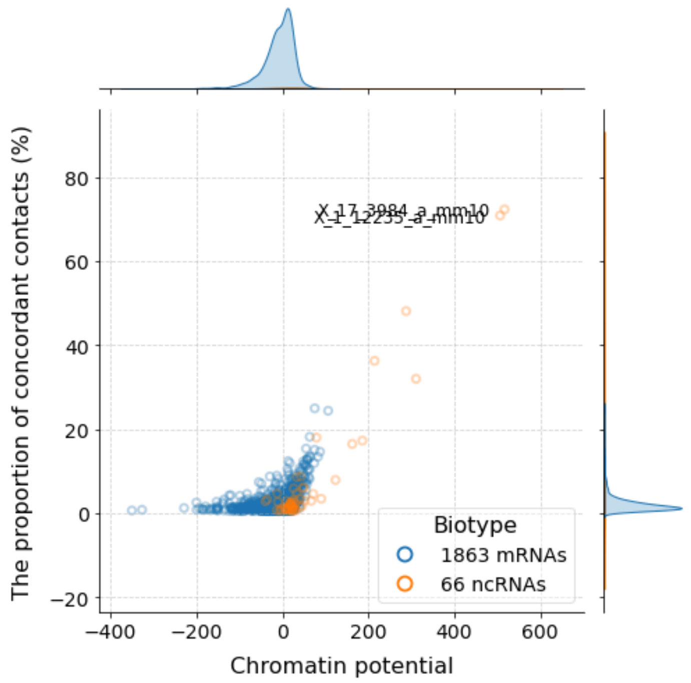


**A2.** RADICL (1% FA), ES, M.mus, all contacts

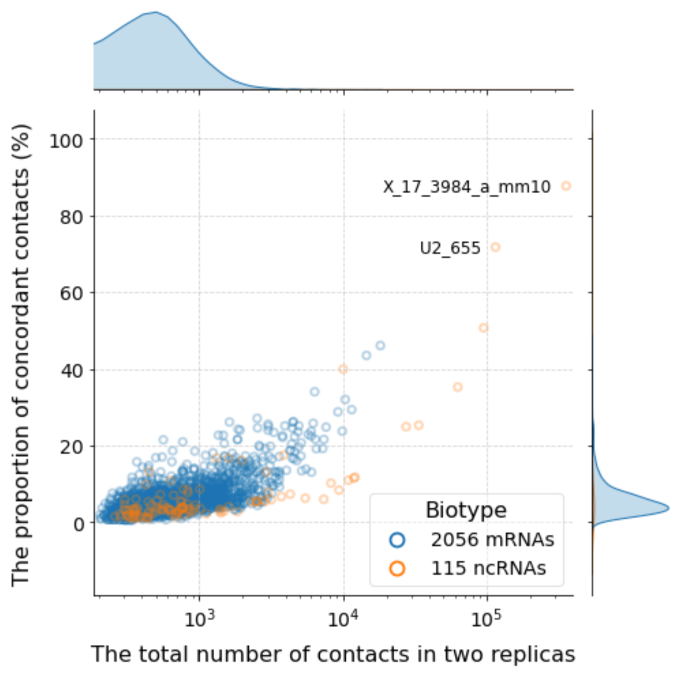


**A3.** RADICL (1% FA), ES, M.mus, contacts from peaks

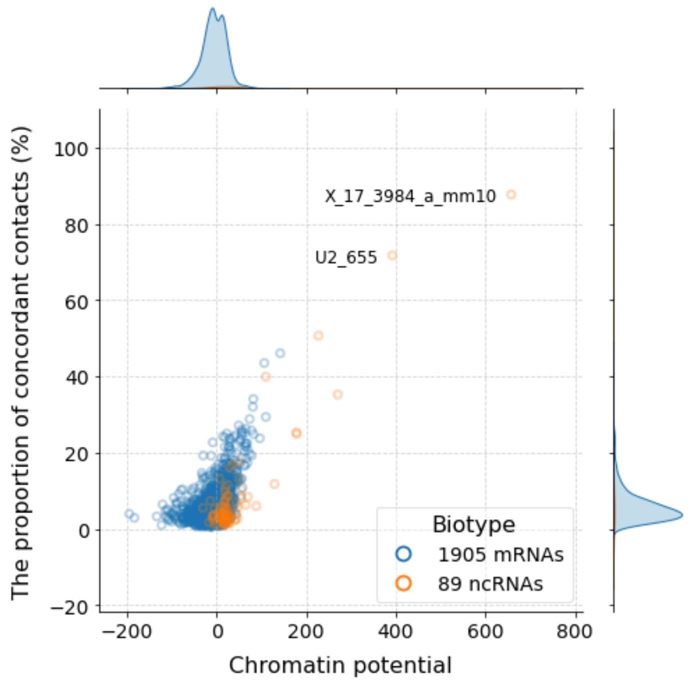


**A4.** RADICL (1% FA), ES, M.mus, contacts from peaks

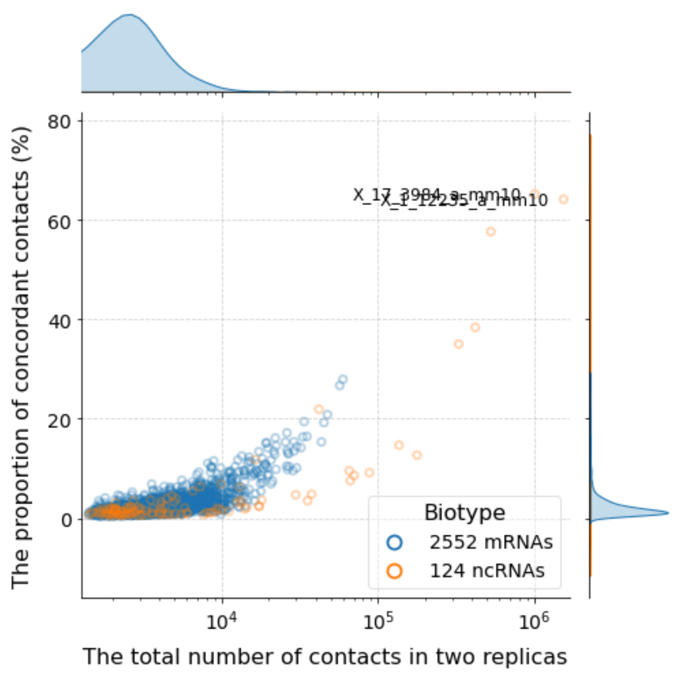


**B1.** RADICL (2% FA), ES, M.mus, all contacts

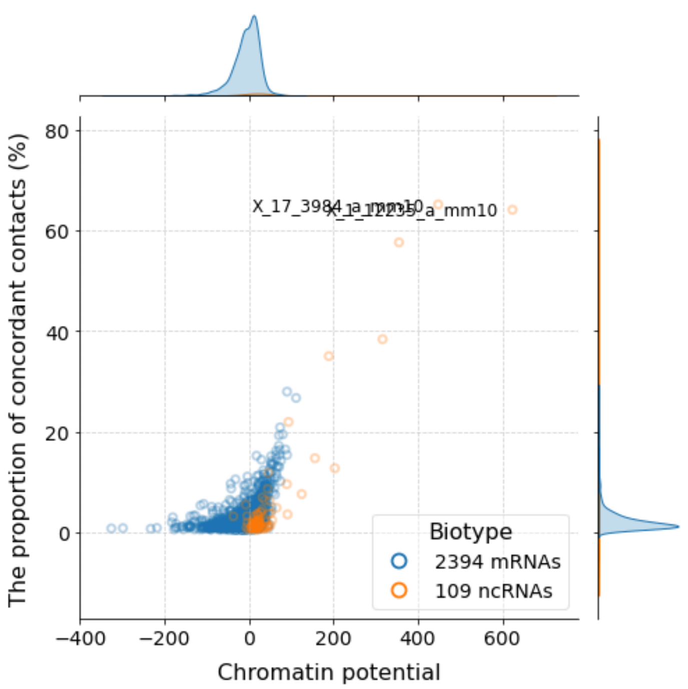


**B2.** RADICL (2% FA), ES, M.mus, all contacts

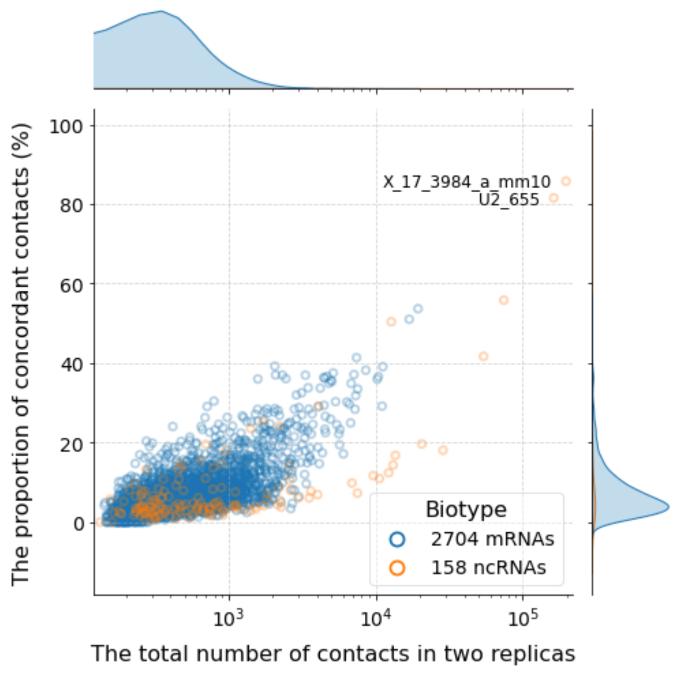


**B3.** RADICL (2% FA), ES, M.mus, contacts from peaks

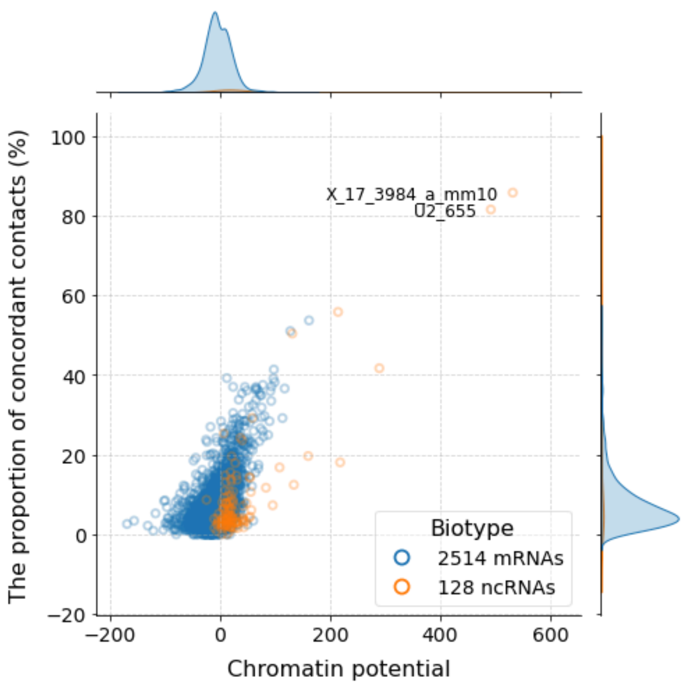


**B4.** RADICL (2% FA), ES, M.mus, contacts from peaks

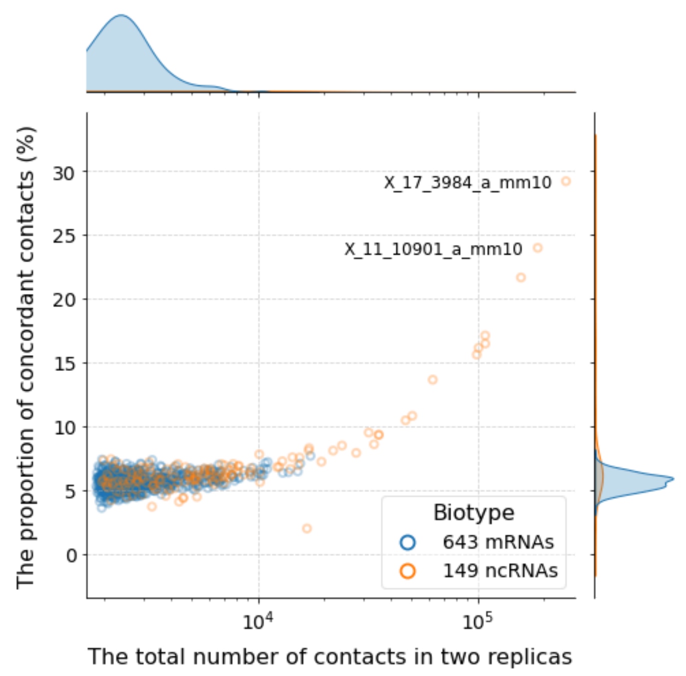


**C1.** RADICL (NPM), ES, M.mus, all contacts

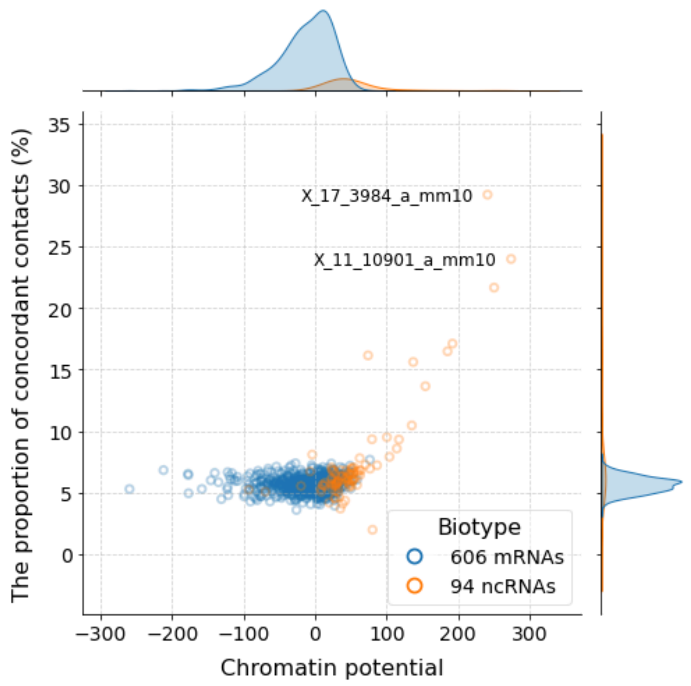


**C2.** RADICL (NPM), ES, M.mus, all contacts

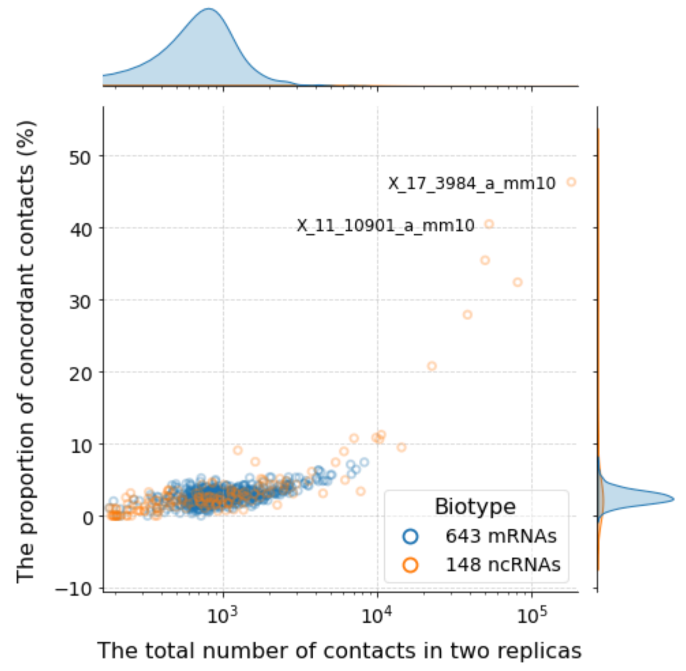


**C3.** RADICL (NPM), ES, M.mus, contacts from peaks

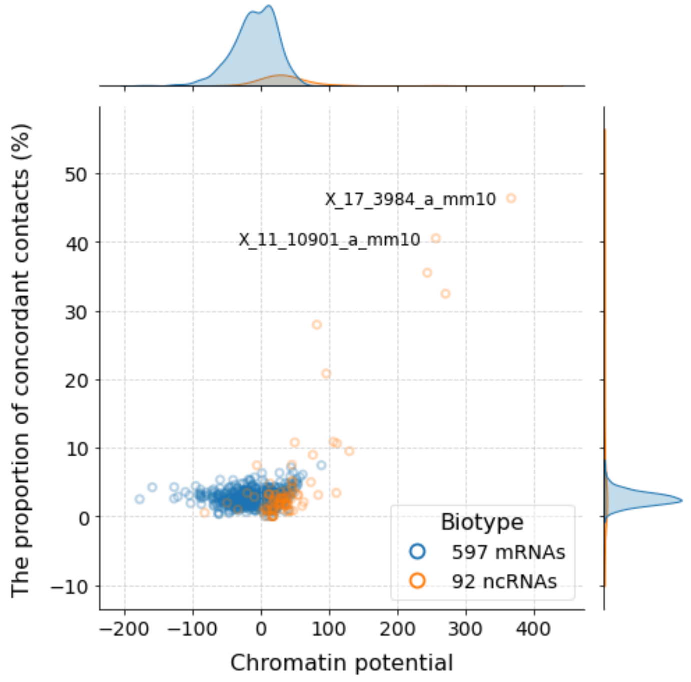


**C4.** RADICL (NPM), ES, M.mus, contacts from peaks

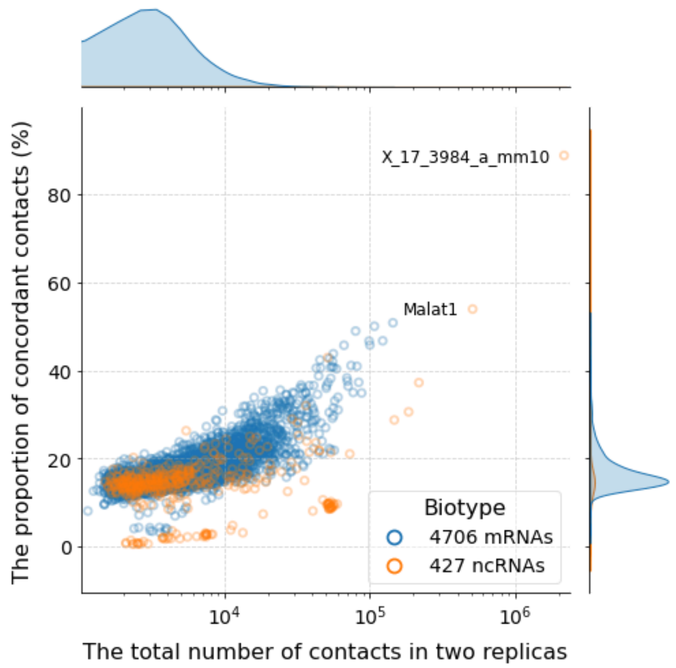


**D1.** GRID, ES, M.mus, all contacts

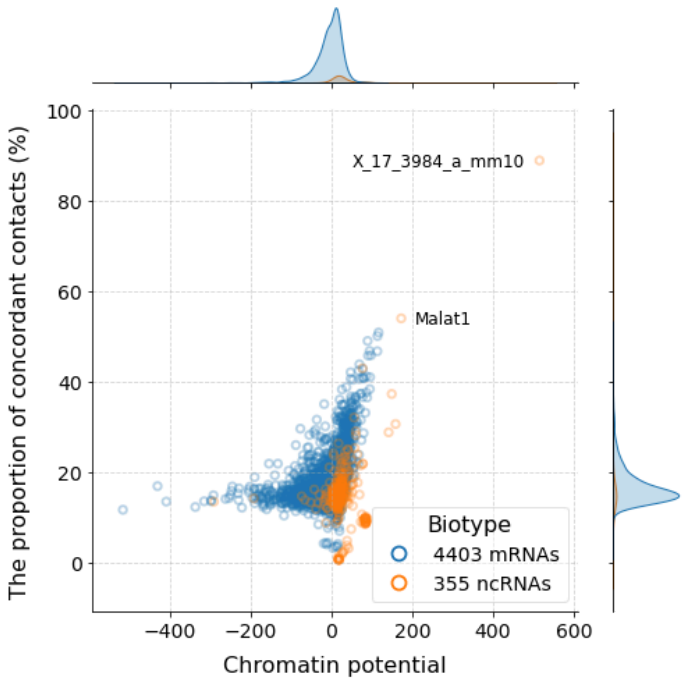


**D2.** GRID, ES, M.mus, all contacts

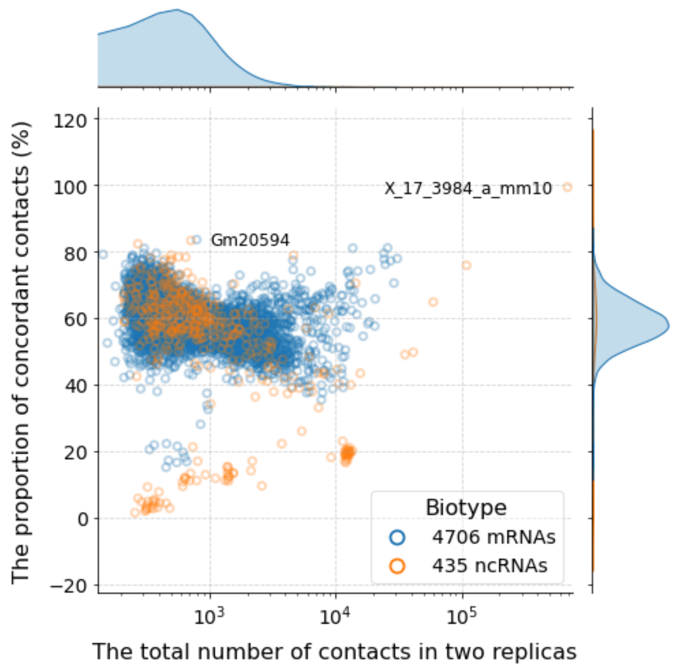


**D3.** GRID, ES, M.mus, contacts from peaks

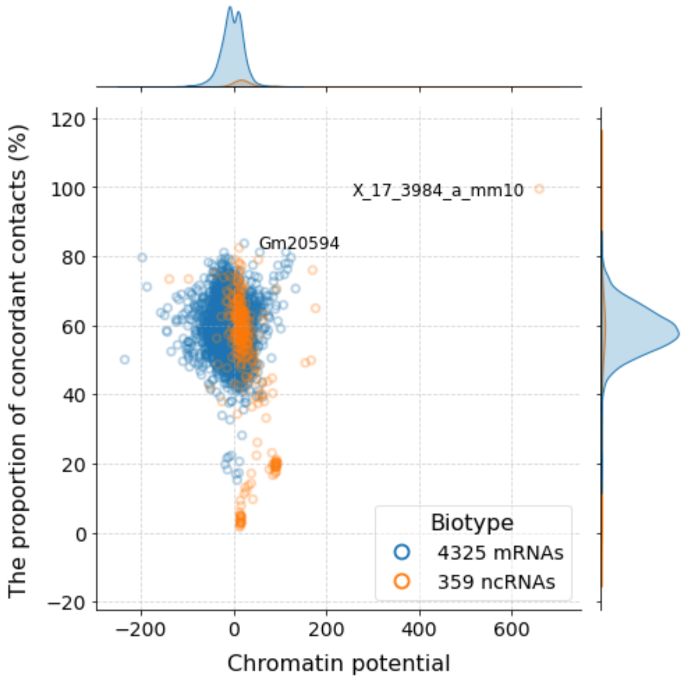


**D4.** GRID, ES, M.mus, contacts from peaks

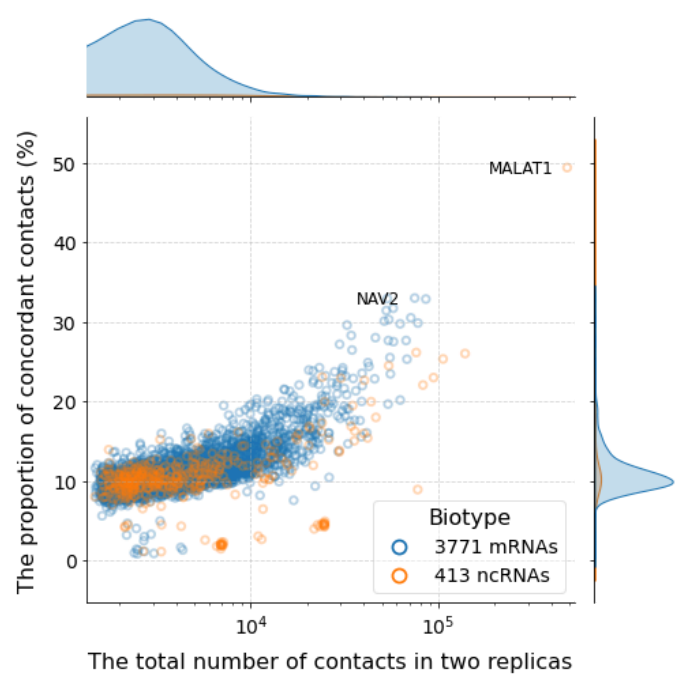


**E1.** GRID, MM.1S, H.sap, all contacts

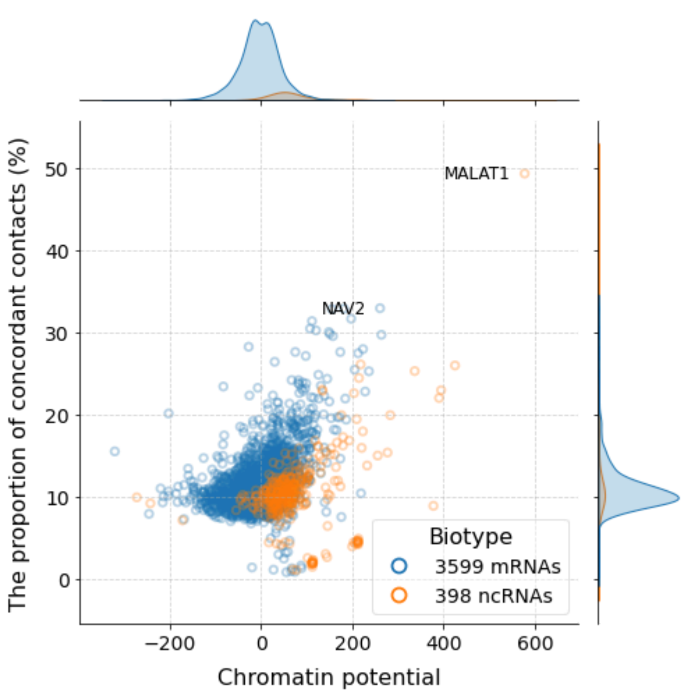


**E2.** GRID, MM.1S, H.sap, all contacts

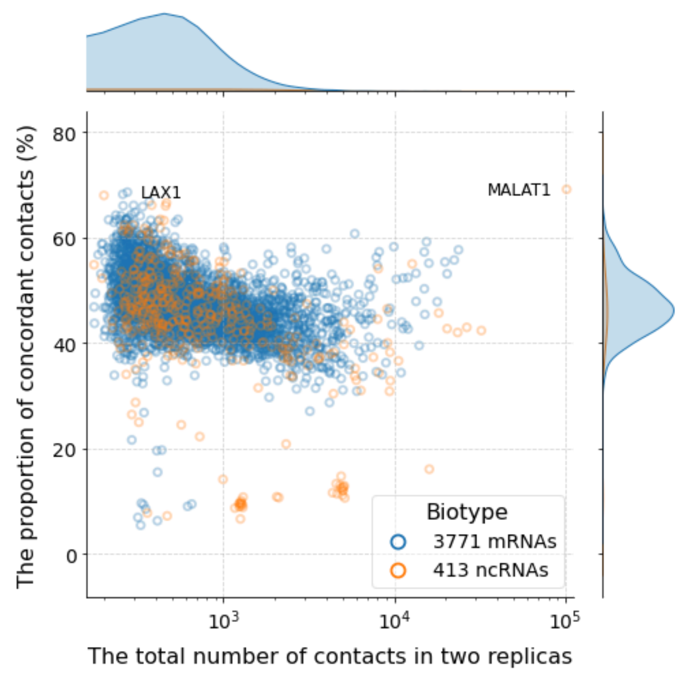


**E3.** GRID, MM.1S, H.sap, contacts from peaks

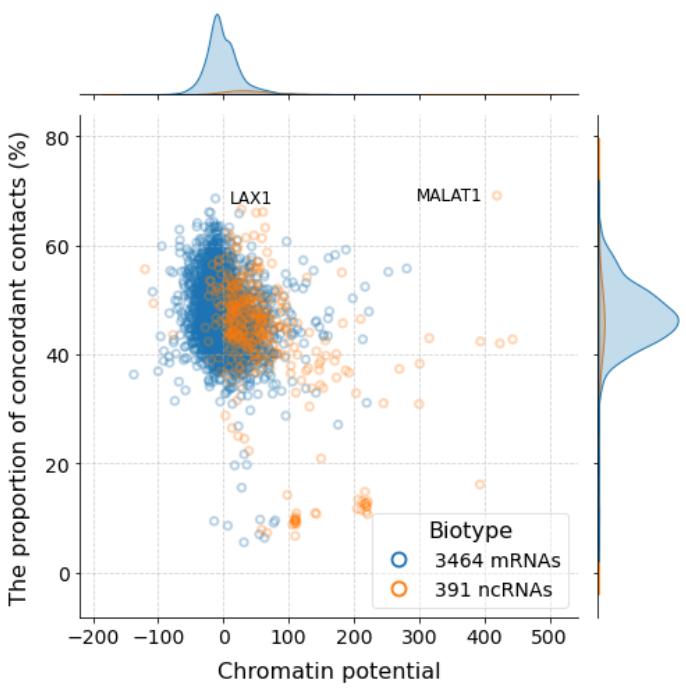


**E4.** GRID, MM.1S, H.sap, contacts from peaks

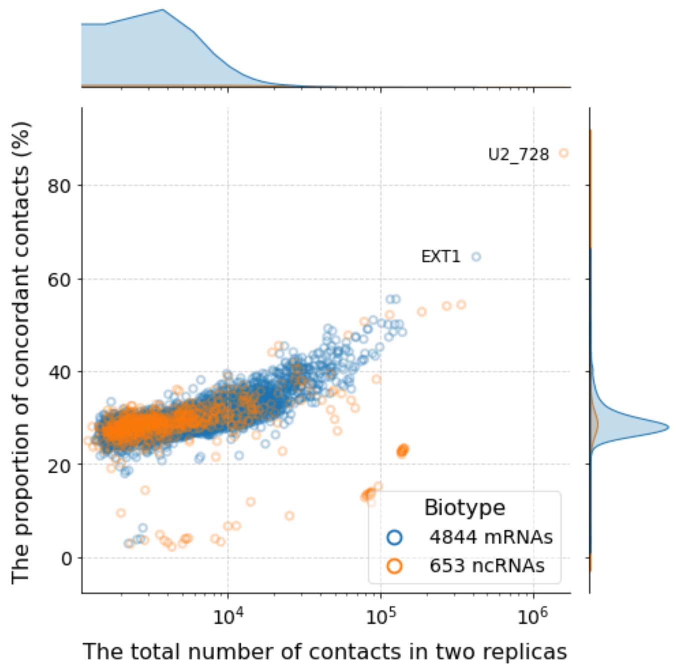


**F1.** GRID, MDA_MB_231, H.sap, all contacts

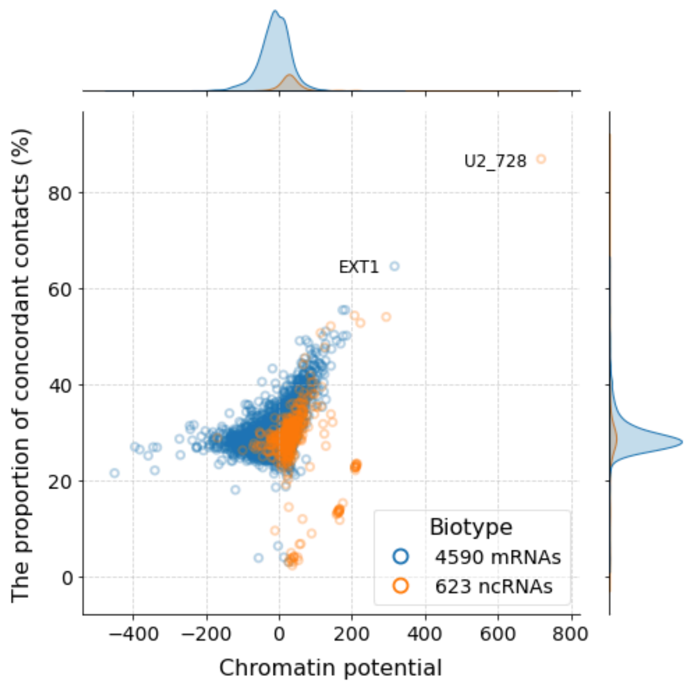


**F2.** GRID, MDA_MB_231, H.sap, all contacts

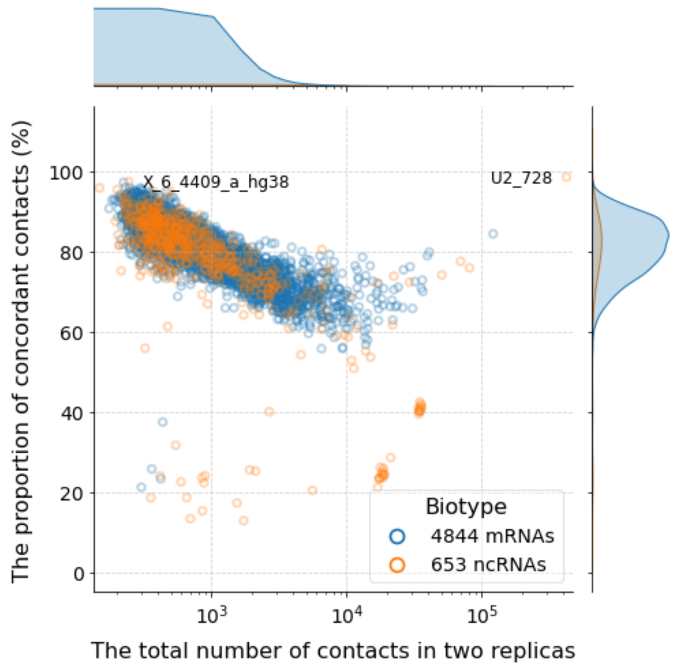


**F3.** GRID, MDA_MB_231, H.sap, contacts from peaks

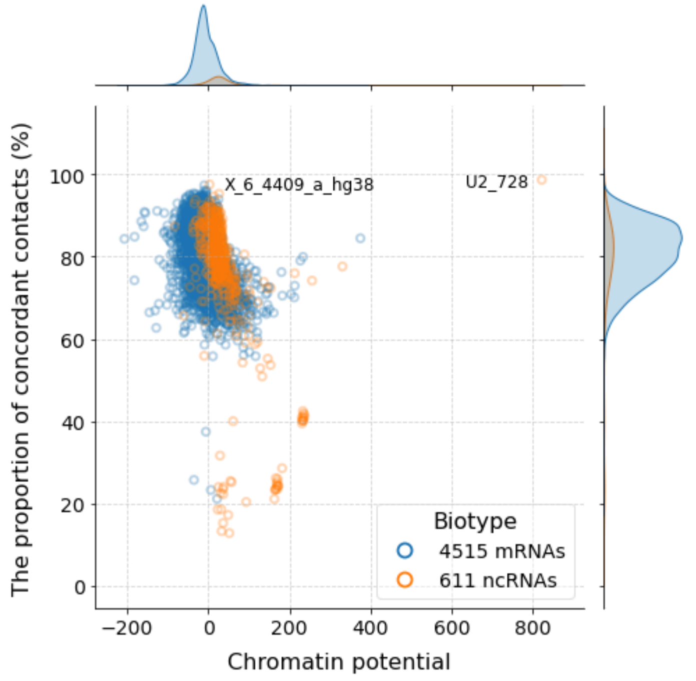


**F4.** GRID, MDA_MB_231, H.sap, contacts from peaks

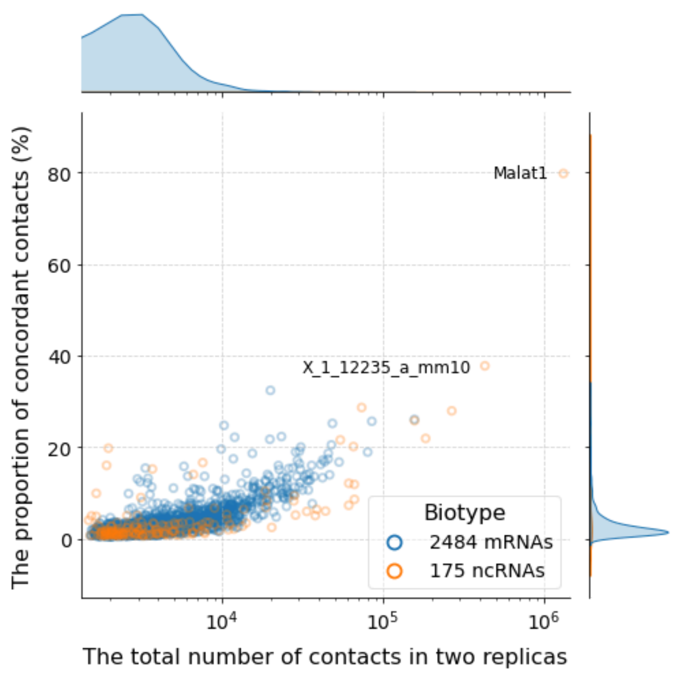


**G1.** RADICL, OPC, M.mus, all contacts

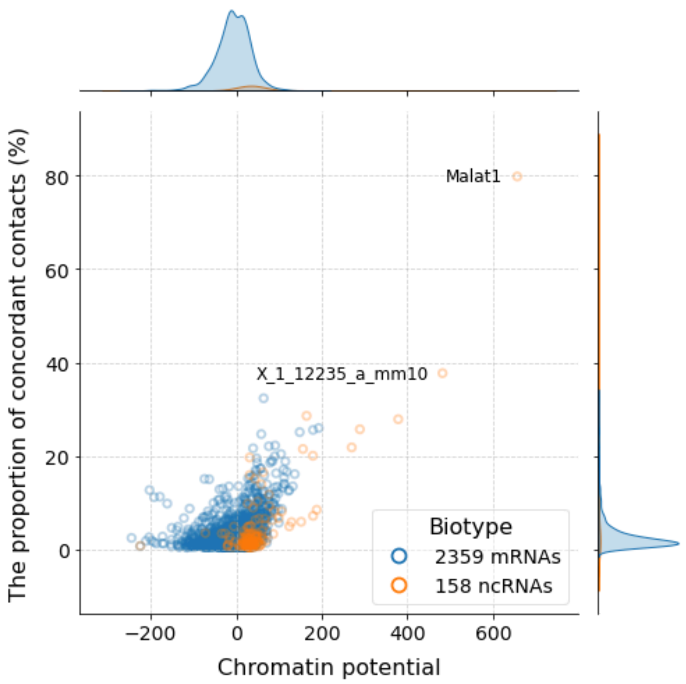


**G2.** RADICL, OPC, M.mus, all contacts

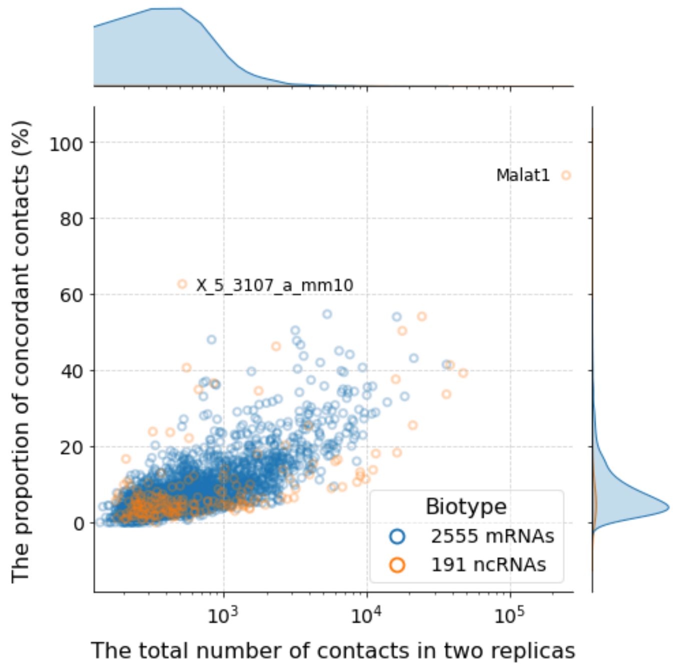


**G3.** RADICL, OPC, M.mus, contacts from peaks

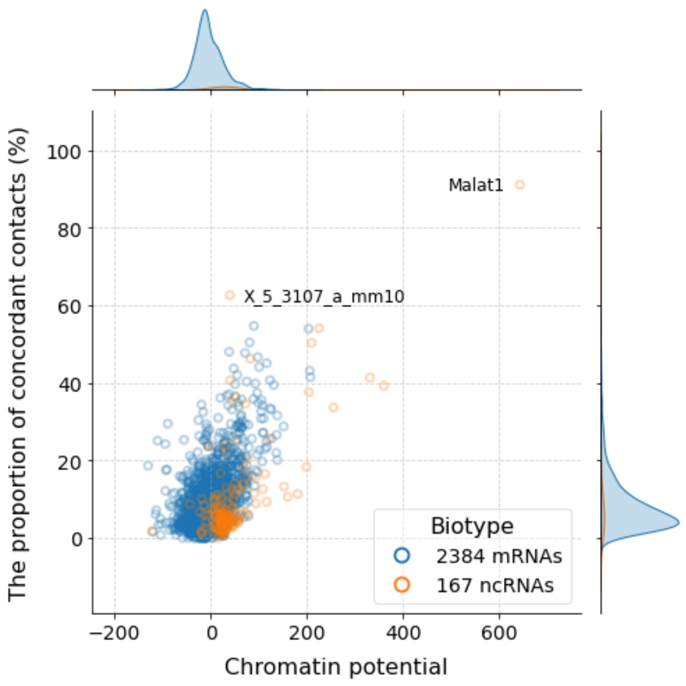


**G4.** RADICL, OPC, M.mus, contacts from peaks

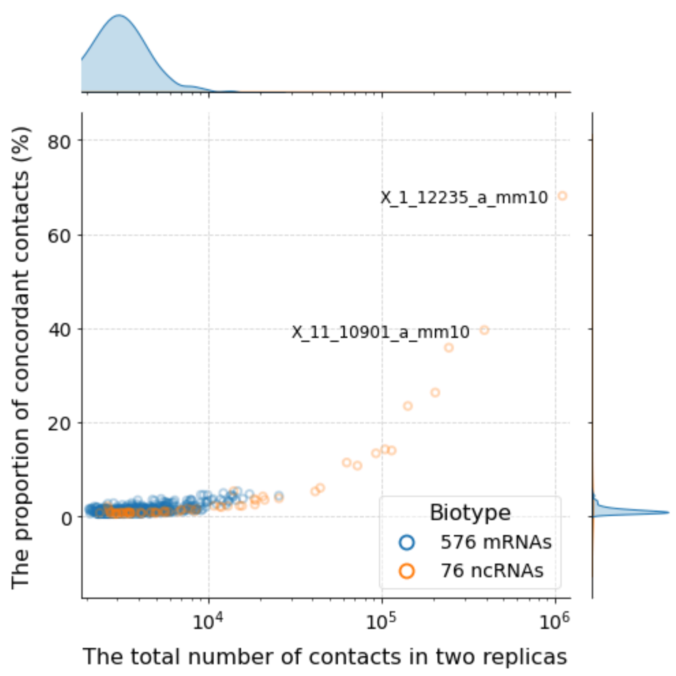


**H1.** RADICL (ActD), ES, M.mus, all contacts

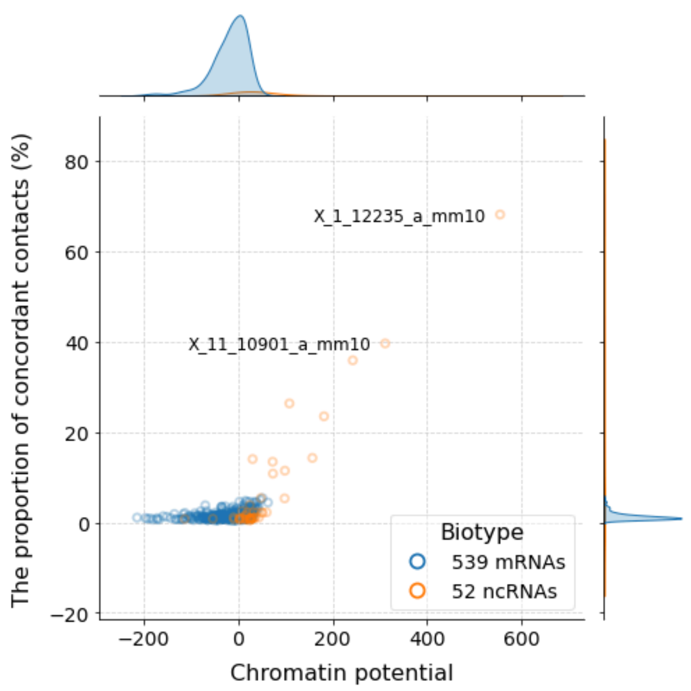


**H2.** RADICL (ActD), ES, M.mus, all contacts

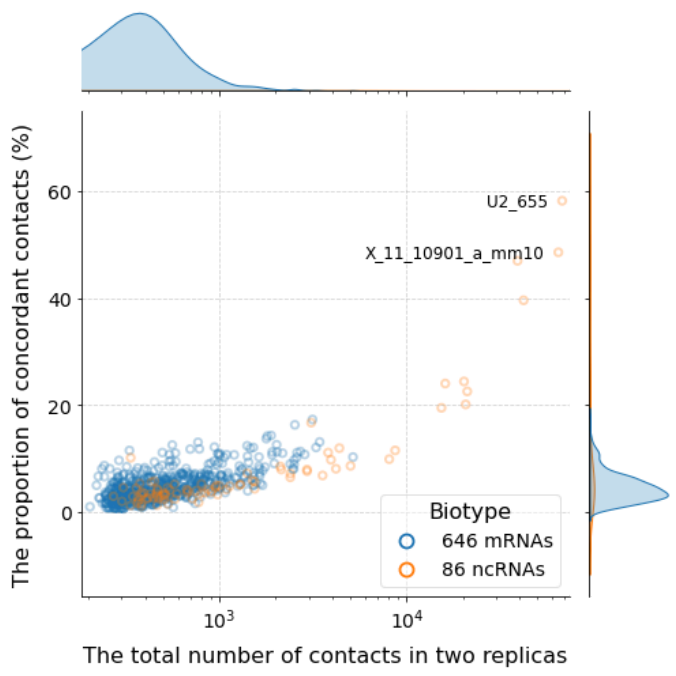


**H3.** RADICL (ActD), ES, M.mus, contacts from peaks

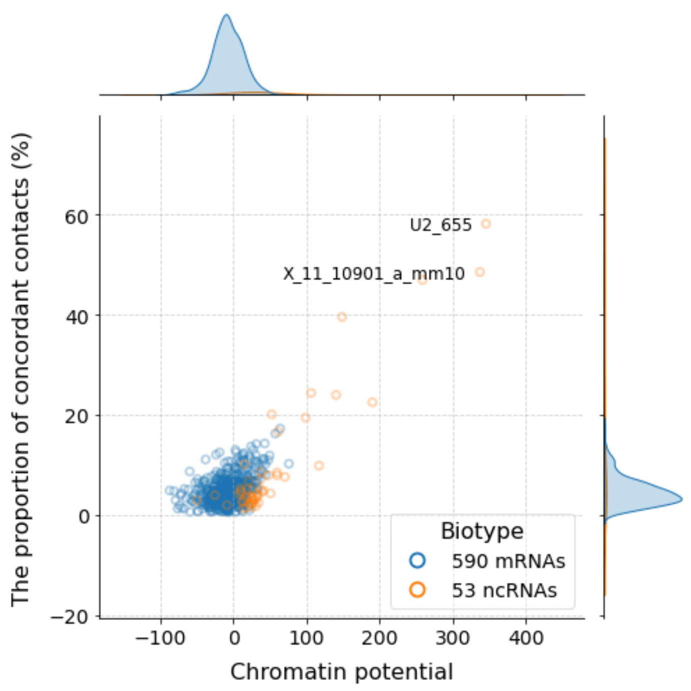


**H4.** RADICL (ActD), ES, M.mus, contacts from peaks

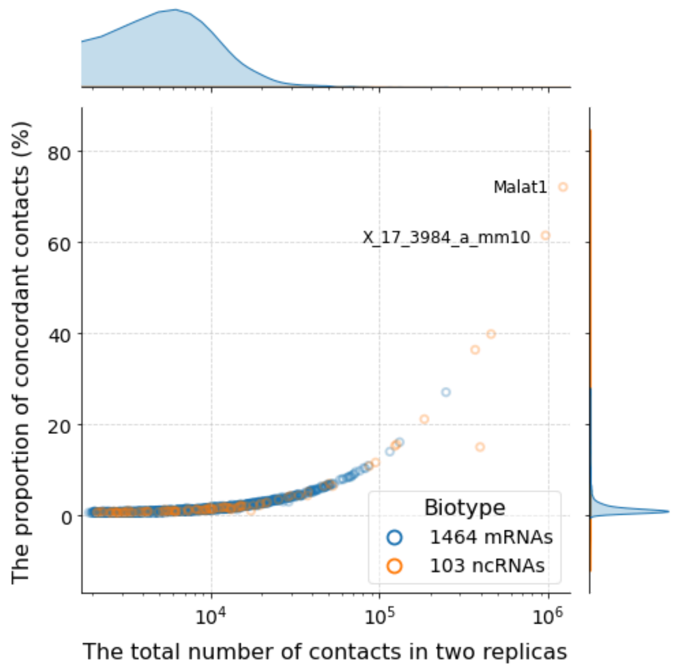


**I1.** RADICL (NPM), OPC, M.mus, all contacts

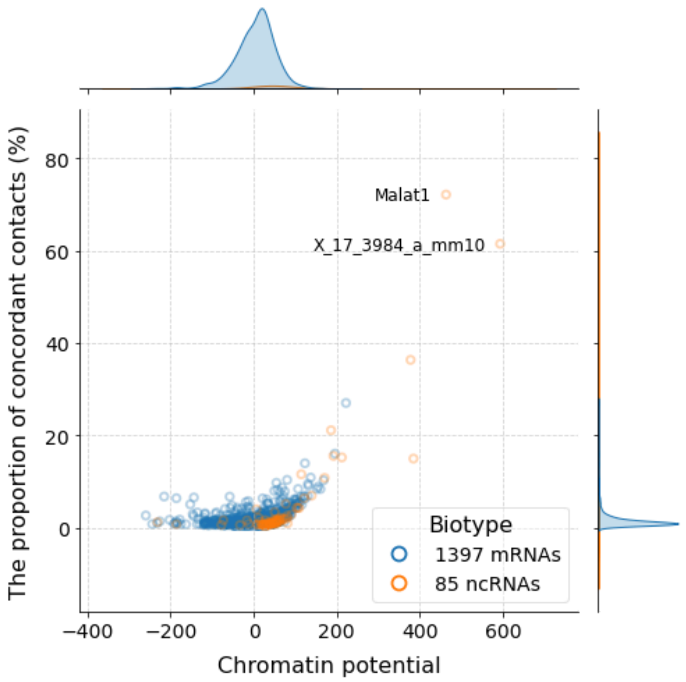


**I2.** RADICL (NPM), OPC, M.mus, all contacts

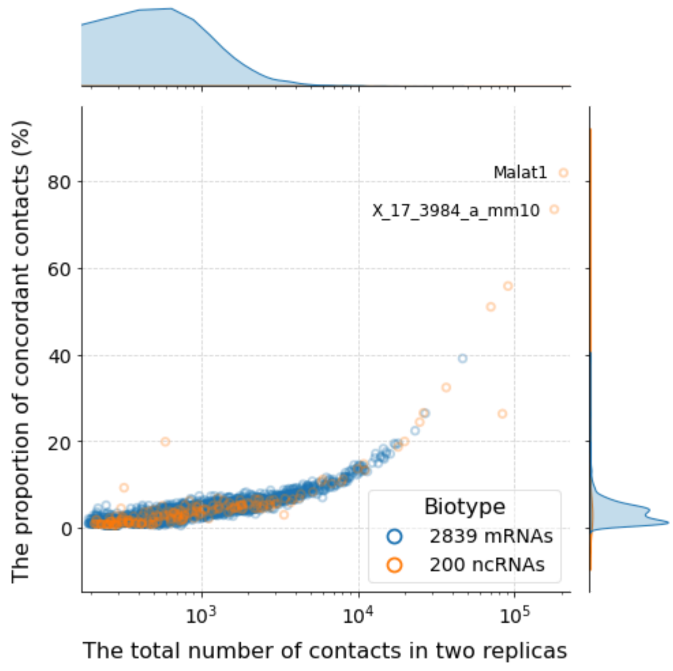


**I3.** RADICL (NPM), OPC, M.mus, contacts from peaks

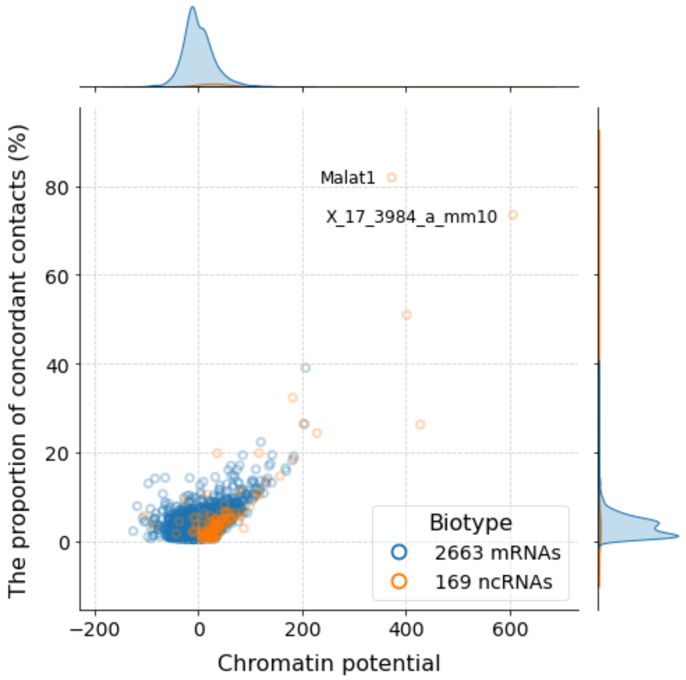


**I4.** RADICL (NPM), OPC, M.mus, contacts from peaks

**Supplementary Figure 3.** Dependence of the proportion of concordant contacts on the total number of contacts, on contacts from peaks and on the chromatin potential for different ATA experiments.

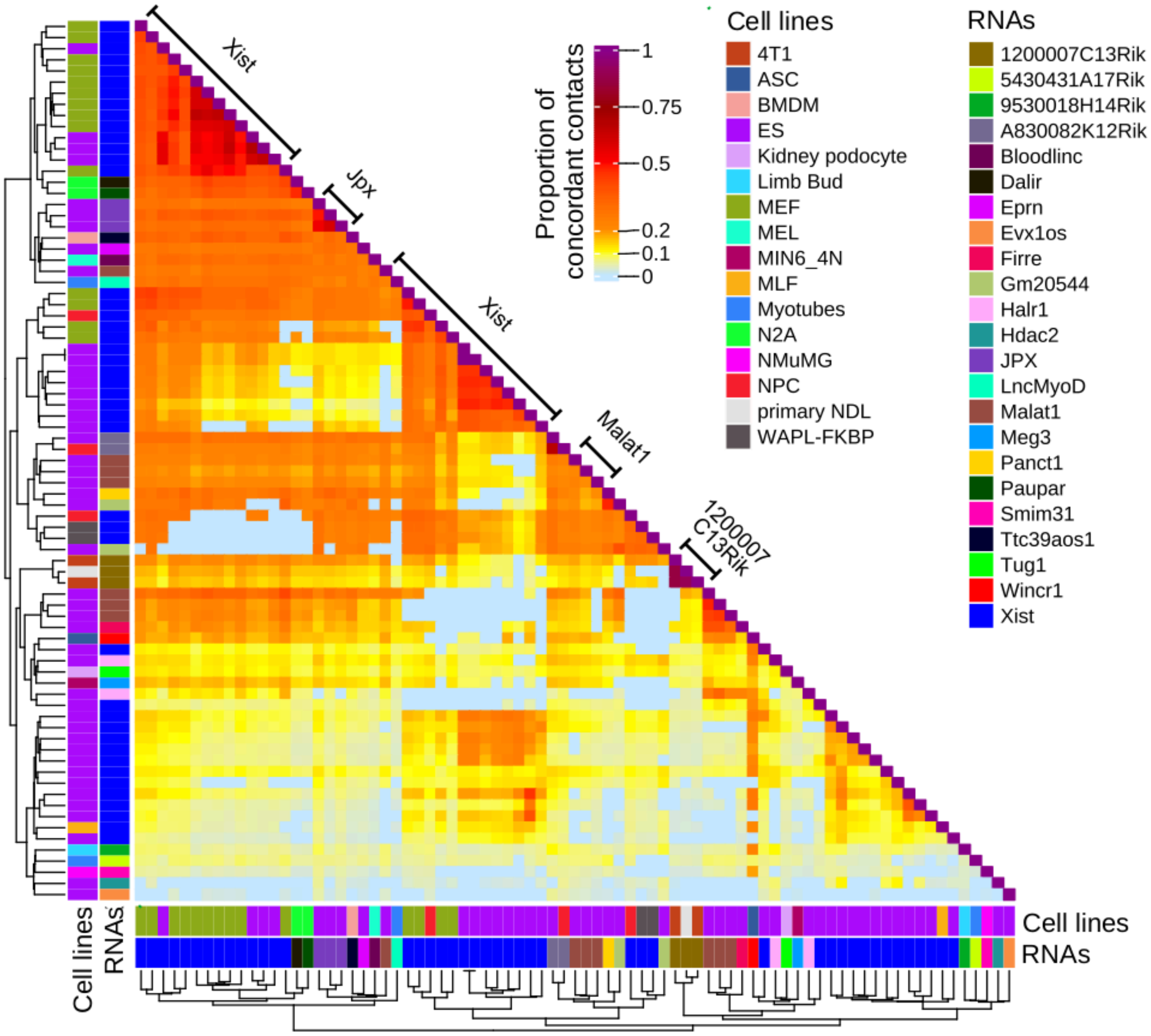


**Supplementary Figure 4.** Heat map showing the proportion of concordant contacts (from BaRDIC peaks) from «one-to-all» experiments for mouse cell lines. Non-significant enrichments (p-value > 0.05) are set to zero. Clustering is performed by cell types and RNAs used in the experiment. The bin size is 1000 nucleotides.
